# A comprehensive map of hotspots of de novo telomere addition in *Saccharomyces cerevisiae*

**DOI:** 10.1101/2023.03.20.533556

**Authors:** Katrina Ngo, Tristen H. Gittens, David I. Gonzalez, E. Anne Hatmaker, Simcha Plotkin, Mason Engle, Geofrey A. Friedman, Melissa Goldin, Remington E. Hoerr, Brandt F. Eichman, Antonis Rokas, Mary Lauren Benton, Katherine L. Friedman

**Affiliations:** Department of Biological Sciences, Vanderbilt University, 465 21st Avenue S, 1210 MRBIII, Nashville, TN 37232; Evolutionary Studies Initiative, Vanderbilt University, 465 21st Avenue S, 1210 MRBIII, Nashville, TN 37232; Department of Biochemistry, Vanderbilt University, 465 21st Avenue S, 1210 MRBIII, Nashville, TN 37232; Department of Computer Science, Baylor University, One Bear Place #97141, Waco, TX, 76798

**Keywords:** de novo telomere addition, DNA repair, DNA damage, genomic instability, yeast, *Saccharomyces cerevisiae*

## Abstract

Telomere healing occurs when telomerase, normally restricted to chromosome ends, acts upon a double-strand break to create a new, functional telomere. De novo telomere addition on the centromere-proximal side of a break truncates the chromosome but, by blocking resection, may allow the cell to survive an otherwise lethal event. We previously identified several sequences in the baker’s yeast, *Saccharomyces cerevisiae*, that act as hotspots of de novo telomere addition (termed Sites of Repair-associated Telomere Addition or SiRTAs), but the distribution and functional relevance of SiRTAs is unclear. Here, we describe a high-throughput sequencing method to measure the frequency and location of telomere addition within sequences of interest. Combining this methodology with a computational algorithm that identifies SiRTA sequence motifs, we generate the first comprehensive map of telomere-addition hotspots in yeast. Putative SiRTAs are strongly enriched in subtelomeric regions where they may facilitate formation of a new telomere following catastrophic telomere loss. In contrast, outside of subtelomeres, the distribution and orientation of SiRTAs appears random. Since truncating the chromosome at most SiRTAs would be lethal, this observation argues against selection for these sequences as sites of telomere addition per se. We find, however, that sequences predicted to function as SiRTAs are significantly more prevalent across the genome than expected by chance. Sequences identified by the algorithm bind the telomeric protein Cdc13, raising the possibility that association of Cdc13 with single-stranded regions generated during the response to DNA damage may facilitate DNA repair more generally.

## Introduction

The maintenance of DNA integrity is essential for cell function. To maintain genomic integrity and prevent sequence loss, most eukaryotic chromosomes terminate with nucleoprotein structures termed telomeres that protect chromosomes from end-to-end fusion and block excessive nucleolytic resection. Telomeres contain a characteristic, repetitive sequence that is rich in thymine and guanine (TG-rich) on one strand (Blackburn 1991). While the majority of the telomere is double-stranded, the TG-rich strand extends past the complementary cytosine and adenine (CA)-rich strand to create a 3’ overhang. Regeneration of this 3’ overhang after each round of DNA replication results in progressive sequence loss, but in cells that maintain telomere length over successive generations, this end-replication problem is counterbalanced through extension of the 3’ strand by telomerase (reviewed in Lingner et al. 1995; Osterhage & Friedman, 2009; Bonnell et al. 2021). Telomerase uses an intrinsic RNA molecule as template for synthesis of the TG-rich strand (Greider & Blackburn, 1989; Singer & Gottschling, 1994) while the lagging strand polymerase machinery subsequently fills in the complementary, CA-rich strand (reviewed in Gilson & Géli, 2007; Pfeiffer & Lingner, 2013).

Because telomeres are, by definition, the end of a DNA molecule, they resemble a DNA double-strand break (DSB). Indeed, similar to telomeres, enzymatic resection at a DSB generates 3’ overhangs that can serve as substrates for homologous recombination. The specific sequence of the 3’ overhang at telomeres distinguishes it from 3’ overhangs generated by resection at a double strand break, thereby enforcing different outcomes at these otherwise similar structures (telomere elongation versus DNA repair, respectively; reviewed in Casari et al. 2022; Doksani & de Lange, 2014). However, rarely, the 3’ overhang generated at a DSB is recognized by telomerase, resulting in addition of a new or de novo telomere (reviewed in Hoerr et al. 2021; Pennaneach et al. 2006). De novo telomere addition (dnTA), also termed telomere healing, causes loss of sequences distal to the site at which the telomere is added but prevents additional resection that would ultimately be lethal.

Several human diseases (e.g. Phelan/Mcdermid syndrome and α-thalassemia) are associated with terminal truncations generated by dnTA (Bonaglia et al. 2011; Guilherme et al. 2015; Lamb et al. 1993; Nevado et al. 2022). The observation of recurrent telomere addition events within a small chromosome region suggests that sequences associated with these diseases are unusually prone to telomerase action. This phenomenon is not limited to human cells, and has been observed in other eukaryotic organisms including *S. cerevisiae* (Mangahas et al., 2001; Ouenzar et al., 2017; Stellwagen et al., 2003). Although dnTA events are generally very rare, the *S. cerevisiae* genome contains hotspots where dnTA occurs at frequencies estimated to be at least 200-fold above background (Obodo et al. 2016; Epum et al. 2020). These sequences present a unique opportunity to use yeast as a model to study the consequences of such sequences for genome stability and evolution.

Telomeres in *S. cerevisiae* have a 3’ terminating strand that consists of irregular repeats containing a pattern of a single T followed by one, two, or three Gs. Despite this heterogeneity, the telomere contains recognition sites for several sequence-specific DNA binding proteins that associate with the double-stranded portion of the telomere and the single-stranded TG-rich overhang (Rap1 and Cdc13, respectively; reviewed in Wellinger & Zakian, 2012). Rap1 participates in telomere length homeostasis, telomere capping, and formation of telomeric chromatin (Hardy et al. 1992; Kyrion et al. 1993; Marcand et al. 1997; Negrini et al. 2007; Pardo & Marcand, 2005; Teixeira et al. 2004; Vodenicharov et al. 2010), while Cdc13 interacts with the Est1 component of telomerase to recruit telomerase to telomeres (Evans & Lundblad, 1999; Pennock et al. 2001, Chen et al. 2018). Cdc13 additionally interacts with Stn1 and Ten1 to limit nucleolytic resection and promote fill-in synthesis by the lagging strand polymerase machinery (Pennock et al 2001; Lin et al. 2021).

Sequences serving as hotspots of dnTA in yeast were first observed as sites of telomere healing in response to an induced DSB on chromosome VII (Mangahas et al. 2001). Subsequently, spontaneous truncations of chromosome V occurring as a result of dnTA were shown to cluster in a small chromosomal region (Myung et al. 2001; Stellwagen et al. 2003; Pennaneach et al. 2006). Following structure/function analysis of the sequence on chromosome V and an additional hotspot on chromosome IX, we named these sequences Sites of Repair-associated Telomere Addition or SiRTAs. SiRTAs contain two TG-rich sequence tracts. One tract (the Core) serves as the direct substrate for telomere addition by telomerase while the second tract (the Stim, located 5’ to the Core on the TG-rich strand) is required for high levels of dnTA at the Core sequence (Obodo et al. 2016). The Stim can be functionally replaced with canonical Cdc13 binding sites or with a sequence designed to artificially recruit Cdc13 (Obodo et al. 2016). Together, these observations support a model in which resection of the 5’-terminating strand following a DSB exposes TG-rich sequences on the 3’ overhanging strand that are bound by Cdc13, with subsequent recruitment of telomerase. Telomere addition is favored at SiRTAs even when the initiating break is artificially induced 2-3 kilobases distal to the eventual site of telomere addition, suggesting that SiRTAs stimulate repair rather than serving as fragile sites *per se* (Obodo et al. 2016).

For the SiRTAs described above, the TG-rich sequence is on the same strand that terminates as a TG-rich 3’ overhang at the nearest telomere, a property we refer to as the “TG-orientation.” On the left arm of a chromosome, SiRTAs in this orientation are TG-rich on the bottom (3’ to 5’ or minus) strand while on the right arm, the TG-rich sequence is on the top (5’ to 3’ or plus) strand. Telomere addition at a SiRTA in the TG-orientation requires a DSB distal to the SiRTA and stabilizes the centromere-containing side of the break. If the SiRTA is distal to all essential genes on that arm (as is true for the SiRTAs on chromosomes V and IX), the resulting terminal deletion is compatible with viability, even in a haploid strain. However, not all characterized SiRTAs are TG-oriented. We recently described a SiRTA in the opposite or “CA-orientation” that promotes cell survival under sulfate-limiting conditions by facilitating formation of an acentric fragment containing *SUL1*, encoding the primary sulfate transporter (Hoerr et al. 2023). Despite identification of several SiRTAs in addition to those described above (Ngo et al. 2020), understanding of the genome-wide frequency and distribution of these sequences is lacking.

Here, we validate the use of a computational algorithm, the <u>C</u>omputational <u>A</u>lgorithm for <u>T</u>elomere <u>H</u>otspot Identification (CATHI), to predict SiRTA function based on similarity with the TG_1-3_ pattern of the yeast telomeric repeat. In parallel, we develop and validate a high-throughput sequencing method that dramatically increases the number of putative sequences that can be characterized while simultaneously yielding information about the site of telomerase action. Together, we use these approaches to determine the overall locations and orientations of SiRTAs on a genome-wide scale. All but one of the subtelomeric repetitive regions (defined as X and Y’ elements; Louis et al. 1994; Louis & Haber, 1992) contain at least one SiRTA in the TG-orientation. However, outside of the subtelomeric regions, there is no apparent bias in the location or orientation of predicted SiRTAs, although these sequences occur more frequently than expected by chance. SiRTA function correlates with the ability of a sequence to bind Cdc13, but overall binding affinity is insufficient to explain all variation in the frequency of dnTA. This work provides a foundation for developing a fuller understanding of how sites with a propensity to stimulate dnTA impact genomic stability and evolution.

## Methods

### Strain construction

Strains were constructed in the S288C background as described (Ngo et al. 2020; Hoerr et al. 2023). The parental strain contains a *URA3* marker distal to an HO recognition site on chromosome VII (YKF1975 *MAT**a**::ΔHOcs::hisG hmlαΔ::hisG HMRa::NAT ura3Δ851 trp1Δ63 leu2Δ::KAN^R^ ade3::GAL10::HO* Chr VII, 15828-16027 (*adh4*)::HOcs::*HYG^R^ pau11*::*URA3*). 300 bp sequences to be tested for SiRTA function were inserted using the CRISPR/Cas9 system as described in Anand et al. (2017) using a guide sequence of 5’-TGCGGCAAGTTCATCTTCCA located ∼2kb centromere-proximal to the HO recognition site. PCR products for recombinational insertion were generated as follows. Forward primers were designed by including 40 bases upstream of the gRNA recognition site (5’TTTCTTTGGAAAACGTTGAAAATGAGGTTCTATGATCTAC) followed by the first ∼20 bases on the 5’ end of the sequence of interest. Reverse primers were constructed by taking the reverse complement of the 40 bases downstream of the gRNA site (5’-AGAACATAGAATAAATTTGGTACTGGAACGTTGATTAACT) followed by the last ∼20 bases of the sequence of interest. Sequences tested are listed in Supplementary File 1. The DNA fragment needed to insert SiRTA 6R210(+) onto chromosome VII using the CRISPR system was synthesized and inserted into the pMX plasmid using Invitrogen GeneArt Gene Synthesis services (Thermo Fisher Scientific). The 6R210(+) DNA fragment was amplified from the plasmid using PCR designed as described above. One step gene replacement using template DNA from pFA6a-TRP1 (Longtine et al. 1998) was used to replace *RAD52*.

For testing on chromosome IX, the *URA3* marker and HO cleavage site were integrated on chromosome IX to create strain YKF1752 (*MAT**a**::ΔHOcs::hisG hmlαΔ::hisG HMRa::NAT ura3Δ851 trp1Δ63 leu2Δ::KAN^R^ ade3::GAL10::HO* Chr9;35050-41450::HOcs::HPH^R^ *soa1*::*URA3*) as described in Obodo et al. (2016). Yeast strains containing the BS Mut1 and BS Mut2 mutations are described in Obodo et al. (2016) as YFK1610 and YFK1652, respectively.

### HO cleavage assay

The HO cleavage assay was performed as described (Ngo et al. 2020; Hoerr et al. 2023). Briefly, cells were grown in synthetic dropout media lacking uracil (SD-Ura) + 2% raffinose to an optical density at 600nm (OD600) of 0.6-1.0. Cells were serially diluted and plated on yeast extract peptone medium with either 2% dextrose (YEPD) or 2% galactose (YEPgal). After incubation at 30°C for three days, colony number was determined on at least two plates of each condition. The frequency of survival on YEPgal was calculated as: (average number of colonies per plate on YEPgal x dilution factor)/(average number of colonies per plate on YEPD x dilution factor). At least 100 colonies surviving on YEPgal were patched to medium containing 1 mg/mL 5-fluoroorotic acid (5-FOA) to select for cells in which the *URA3* marker was lost [gross chromosomal rearrangement (GCR) events]. The frequency of GCR events was determined as: (frequency of survival on YEPgal*frequency of clones demonstrating 5-FOA resistance). Thirty clones that displayed growth on medium containing 5-FOA were selected and inoculated in liquid YEPD for genomic DNA extraction using the MasterPure™ Yeast DNA Purification Kit (Lucigen). Multiplex PCR was used as described in Ngo et al. (2020) to map the approximate site of dnTA in relationship to the sequence of interest. Primers for chromosome VII and IX are listed in Supplementary File 1. Colonies where the DNA loss event mapped within the sequence of interest were tested for telomere addition using one primer centromere proximal to the putative telomere addition site and a second primer complementary to the telomeric repeat (Supplementary File 1).

### Pooled Tel-seq

Thirty 5-FOA resistant clones isolated as described above were separately inoculated in 200 µL of YEPD in a 96-well culture plate and incubated overnight at 30°C to reach saturation. Equal volumes (at least 30 µL) of each culture were pooled and DNA was extracted using the YeaStar^TM^ genomic DNA kit (ZYMO research).

Libraries were prepared using 50 ng of genomic DNA and a modified protocol using the Twist Library Preparation kit (Twist Bioscience 106543). Denatured DNA templates in a 96-well plate were randomly primed with 5’ barcoded adapters. Samples were pooled, captured on streptavidin coated magnetic beads, and washed to remove excess reactants. A second 5’ adapter tailed primer with a strand-displacing polymerase was utilized to convert the captured templates into dual adapter libraries. Beads were washed to remove excess reactants. Four cycles of PCR were utilized to amplify the library and incorporate the plate barcode in the index read position. Libraries were sequenced using the NovaSeq 6000 with 150 bp paired end reads targeting 13 to 15 million reads per sample. Real Time Analysis software (version 2.4.11; Illumina) was used for base calling and data quality control was completed using MultiQC v1.7.

Sequencing data are available from the NIH Sequence Read Archive (SRA) under BioProject ID <u>PRJNA939836</u>. Reads mapping to a 300 bp control sequence located in the essential gene *BRR6* (Chr VII: 36933 to 37233) or to the 300 bp sequence of interest [inserted on chromosome VII or at the endogenous location of SiRTA 9L44(-)] were identified using Bowtie2 (Galaxy Version 2.5.0+galaxy0) with the sensitive local setting (Langmead et al. 2009; Langmead and Salzberg 2012). Any remaining library primer sequences were removed using the Trimmomatic tool (Galaxy Version 0.38.0; Bolger et al. 2014) and reads mapping to the putative SiRTA that also contain telomere sequence (match to 5’-GGGTGTGG or 5’-CCACACCC) were identified and tabulated. The number of individual reads with evidence of telomere addition was normalized to the number of control reads at *BRR6* by expressing the number of telomere reads as a percentage of control reads. Where applicable, sites of telomere addition were mapped to the original SiRTA sequence to determine the location of the event.

### Purification of Cdc13-DBD

Cdc13-DBD was expressed in *E. coli* using pET21a-Cdc13-DBD-His6, a gift from the Wuttke lab. Purification was done as described (Anderson et al. 2002; Obodo et al. 2016).

### Fluorescence Polarization binding assays

Binding assays were conducted using a fixed concentration of a 5’-6-carboxyfluorescein (FAM) labeled tel-11 oligonucleotide (25 nM). Cdc13-DBD was added at final concentrations of 0, 6.25, 12.5, 25, 37.5, 50, 62.5 and 75 nM. Competition binding assays were conducted at fixed concentrations of Cdc13-DBD (30 nM) and 5’-6-FAM labeled tel-11 oligonucleotide (25 nM). Unlabeled oligonucleotides of 75 bases each were used at final concentrations of 0, 6.25, 12.5, 25, 50, 150 and 200 nM. Oligonucleotide sequences are listed in Supplementary File 1. Labeled and unlabeled oligonucleotides and protein were mixed in binding buffer (50 µM Tris pH 8, 1 µM EDTA pH 8, 15% glycerol, 75 µM NaCl, 75 µM KCl) in a final volume of 80 µL and incubated at 4°C for 30 minutes. Each reaction was measured in triplicate (25 µl per measurement) in a Corning 384 well assay plate using the BioTek Synergy H1 hybrid reader. This procedure was repeated at least three times for each competitor. The relative polarization (ΔP) was determined using the following equation: ΔP=P_0_-P_x_, where P_0_ is the polarization value at a competitor concentration of 0 and P_x_ represents the polarization value at x competitor concentration. For each experiment, technical replicates were averaged and the averaged data were fit to the following equation: ΔP=(P_max_[competitor])/(*K*_i,app_[competitor]), where P_max_ is the maximal polarization value and *K*_i,app_ is the apparent inhibition constant (Anderson et al. 2008; Vaasa et al. 2009). An unlabeled 75mer containing the Tel11 sequence at the center of the oligonucleotide (Tel11-75) was included in each experiment and normalized *K*_i,app_ values are reported as the fold change relative to this control (*K*_i,app_ of Tel11-75/*K*_i,app_ of experimental oligonucleotide)

### Implementation of the CATHI algorithm

Initially, the program generates a series of sliding windows to be utilized in the score calculation. The window and step size of the sliding windows can be customized using the --window and --step options. For each window, the program searches for strings of at least 4 characters that begin with a G and consist of only Gs and Ts. These become the set of candidate scoring regions. From these candidates, regions that consist only of Gs are removed. The program then scans candidate regions for any consecutive Ts, or four or more consecutive Gs. If either are encountered, that candidate region is truncated after the first T or the third G, respectively. Once the set of candidates has been filtered, the number of nucleotides remaining in the candidate set is counted and any applicable scoring penalties are subtracted. There are no penalties applied by default, but users can choose to apply them. The -- penalty option imposes score deductions for any GGTGG sequences, and the --ttpenalty imposes score deductions for Ts that flank the candidate regions.

CATHI is implemented in Python (version 3) using the BioPython (Cock et al. 2009), NumPy (Harris et al. 2020), Pandas (McKinney 2010), and pybedtools (Dale et al. 2011) libraries. CATHI can perform in two modes: (1) score mode; and (2) signal mode. The default score mode will return the maximum CATHI score for each input sequence. For each sequence in the provided FASTA file, CATHI will generate sliding windows and calculate the CATHI score for each window, returning only the maximum score per sequence. In signal mode, CATHI will generate sliding windows and return the genomic coordinates and CATHI score for each window. CATHI output is printed to the screen for easy redirection and can be optionally printed in BED format. Code can be obtained from https://github.com/bentonml/cathi. In this work, both strands of each chromosome in *S. cerevisiae* were separately scanned in signal mode using a step size of 1 and window size of 75. Perfect telomeric repeats representing *bona fide* terminal telomeres were trimmed prior to analysis (if present). Coordinates used for each chromosome are in Supplementary File 2.

Overlapping and adjacent windows meeting or exceeding the threshold value can be merged into a single region using the --cluster option, where the beginning is the start coordinate of the most upstream window and the ending is end coordinate of the most downstream window. The score of the merged region is the maximum CATHI score across all merged windows.

### Generation of a shuffled yeast genome

A set of five randomized versions of the *S. cerevisiae* (sacCer3) genome was generated to evaluate the number of SiRTAs expected when applying the CATHI algorithm to a null model. DNA sequence was downloaded from the sacCer3 reference genome using the BedTools (version 2.30.0) ‘getfasta’ command (Quinlan and Hall 2010) after adjusting the start and end coordinates of each chromosome to exclude subtelomeric regions (Supplementary File 2). Nucleotides were randomly shuffled within each adjusted chromosome using Python’s built-in randomization library. This procedure maintains the nucleotide composition and length of each chromosome while randomizing the actual DNA sequence.

### Enrichment for genomic annotations in putative SiRTAs

Overlap between putative SiRTAs and other genomic annotations was determined using a permutation-based enrichment test. Enrichment for SiRTAs with several different genomic annotations was determined: (1) essential and non-essential genomic regions (Giaever et al. 2002); (2) Est2 binding sites (Pandey et al. 2021); (3) Pif1 binding sites (Paeschke et al. 2011); (4) γH2AX binding sites (Capra et al. 2010); (5) G-quadruplex regions (Capra et al. 2010) and (6) Rap1 binding sites (Rhee and Pugh 2011). When the original dataset included strand information (as in the case of G4-sites) that information was considered in the analysis.

Enrichment between the SiRTAs and the annotations was calculated as the fold change between the observed and expected overlap. To ensure meaningful overlaps between the SiRTAs and the genomic annotations, at least 50% of the binding site was required to overlap with the SiRTA or at least 50% of the SiRTA was required to overlap with the essential/non-essential region. To create the distribution of expected overlap, 1000 permutations were performed by randomly shuffling regions throughout the genome and calculating the amount of SiRTA overlap. Shuffled regions are non-overlapping, length- and strand-matched (G4 sequences only) with the annotations. When specified, telomeric and/or subtelomeric regions were excluded. Subtelomeric regions are defined in Supplementary File 2. For G4 sites, overlap was only recorded if the G4-forming sequence and SiRTA are on the same strand. An empirical p-value is calculated for the overlap using the expected distribution; where relevant, p-values are corrected for multiple comparisons using the Bonferroni method.

### Determining overlap with genes

The location of predicted SiRTAs was compared to the location of genes within the *S. cerevisiae* genome to determine the number of predicted SiRTAs in both inter- and intragenic regions. Coordinates for genes and subtelomeres (defined as X and/or Y’ elements) were obtained from the *S. cerevisiae* S288C annotation available from NCBI (accession numbers NC_001133.9, NC_001134.8, NC_001135.5, NC_001136.10, NC_001137.3, NC_001138.5, NC_001139.9, NC_001140.6, NC_001141.2, NC_001142.9, NC_001143.9, NC_001144.5, NC_001145.3, NC_001146.8, NC_001147.6, NC_001148.4); FASTA and GFF3 files for the reference assembly of strain S288C (GCF_000146045.2) were downloaded from NCBI’s RefSeq database. The RefSeq genome annotation is identical to that in the *Saccharomyces* Genome Database (SGD).

Coordinates for predicted SiRTAs were obtained from the CATHI program and manually converted into GFF3 files, one for each chromosome. Overlap between predicted SiRTAs and genes was calculated using the “intersect” function within bedtools v2.30.0 (Quinlan 2014) for each chromosome. Predicted SiRTAs within annotated genes were manually assigned to the template or coding strand using chromosome visualization in Geneious Prime v2020.1.2.

### Modeling SiRTA distribution

Python programs to model the expected distribution of telomere addition events within a region and to model the random distribution of SiRTAs between the forward and reverse strand are available at https://github.com/geofreyfriedman/sirta. For the latter, random strand distributions were generated for each chromosome based on the observed number of SiRTAs on each strand. The expected distribution of SiRTAs between the forward and reverse strands was quantified by 1) determining the number of consecutive SiRTAs on the same strand (run length) or 2) summing the number of times that consecutive SiRTAs are found on different strands (number of strand switches). To avoid “edge effects” generated at the ends of each chromosome, 10,000 iterations were generated for each chromosome and run lengths or strand switches were summed across the 16 chromosomes (sum of iteration 1, sum of iteration 2, etc). In each case, the observed value was compared to the random distribution generated from 10,000 iterations.

## Results

### High throughput sequencing of pooled samples accurately measures de novo telomere addition

Measurement of the propensity for de novo telomere addition (dnTA) across the genome is complicated by varied chromosome context (which can affect the frequency of competing repair pathways) and our inability to capture dnTA addition events at sequences that are proximal to essential genes and/or in the CA-orientation. To circumvent these limitations, we previously developed a “test site” on the left arm of chromosome VII. CRISPR/Cas9 is used to insert sequences (typically 300 bp) oriented such that the TG-rich sequence of interest is on the bottom (3’ to 5’) strand. A recognition site for the homothallic switching (HO) endonuclease is located ∼2kb distal to the CRISPR/Cas9 integration site. A *URA3* marker located further distal to the HO cleavage site facilitates selection for cells carrying a truncated chromosome VII-L (Figure 1a and b). Importantly, *RAD52* is deleted to prevent homology-directed repair between the sequence inserted on chromosome VII and that same sequence at its endogenous location.

**Figure 1.**
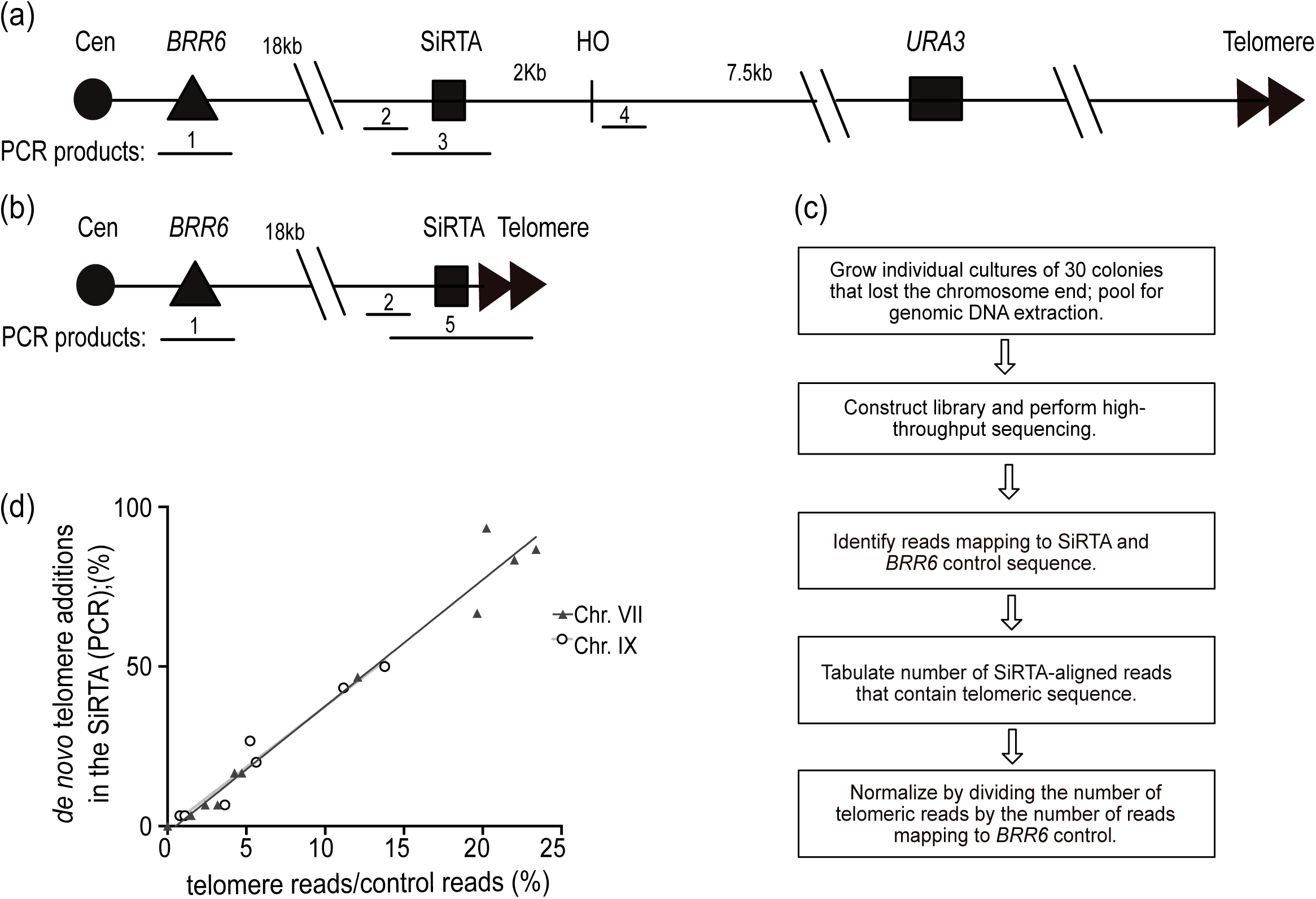
Validation of Pooled Telomere sequencing (PT-seq) as a method to quantify de novo telomere addition. a) Diagram depicting the structure of chromosome VII and the strategy for mapping GCR events resulting from a DSB generated by the HO endonuclease. Locations of the essential gene *BRR6* and the sequence to be tested (SiRTA) are shown. A *URA3* marker inserted distal to the HO cleavage sites allows selection for cells that lose the chromosome terminus following HO cleavage (see text for additional explanation). The approximate locations of PCR products utilized to map GCR events are shown. Product 1 amplifies sequences within *BRR6* and is utilized as a positive control. Products 2 and 3 amplify regions internal to the putative SiRTA or across the SiRTA, respectively, and are used to identify clones in which a GCR event occurred within the SiRTA. Product 4 is used to verify loss of the chromosome terminus. b) Diagram of a GCR event in which telomere addition occurred in the putative SiRTA. Addition of telomeric DNA at the SiRTA is verified by the presence of PCR product 5, generated with a forward primer proximal to the SiRTA and a reverse primer complementary to the telomeric repeat. c) Flow chart representing the PT-seq methodology. D) Correlation of the PCR-based and PT-seq methodologies (Supplementary File 3). Results on chromosome VII (black triangles; r^2^=0.97) and chromosome IX (open circles; r^2^=0.95) were analyzed by linear regression (lines are nearly superimposed). Line equations (VII: y=3.961x-2.039 and IX: y=3.8022x-0.5505) are statistically indistinguishable (p=0.94) by analysis of covariance.

Cells are plated on solid medium containing galactose to induce expression of the HO endonuclease. To escape persistent cleavage and generate a colony, a cell must incur a mutation at the HO site that blocks nuclease recognition or lose the HO site completely through formation of a gross chromosomal rearrangement (GCR). To identify the latter, which include dnTA events, 100 clones arising on the galactose plate are screened for loss of the *URA3* marker via growth on medium containing 5-fluoroorotic acid (5-FOA). Thirty 5-FOA-resistant clones are then analyzed to determine the nature of the resulting GCR event. In past work, we utilized a clone-by-clone mapping strategy that employed multiple PCR reactions to identify the approximate location of each GCR event (Figure 1a). For colonies in which the event maps to the sequence of interest, Southern blotting or PCR utilizing a telomeric primer is utilized to determine if the event involved telomere addition (Figure 1b).

The efficiency of SiRTA function is expressed as the percent of 5-FOA-resistant clones (from a total of 30) that contain a telomere-addition event within the sequence of interest at the insertion site on chromosome VII. Typically, the experiment is done 2-3 times and the average values of the biological replicates are reported. The 300 bp sequence analyzed represents ∼1.4% of the 21,922 bp region within which a GCR event can be recovered [between the HO cleavage site and the first essential gene on VII-L (*BRR6*)]. To determine a threshold for SiRTA activity, we modeled the expectation for random repair within this region (*Materials and Methods* and Supplementary Figure 1). Assuming random distribution of 30 GCR events, two or more would be expected to occur within the 300 bp test sequence in 6.2% of trials, while three or more would be expected in only 0.78% of trials. Therefore, we chose to define a SiRTA as a sequence in which the average efficiency of dnTA is >6.6% (an average of greater than 2 out of 30 clones containing dnTA within the 300 bp test sequence). Sequences tested are named using the following scheme: chromosome number, chromosome arm (L for left and R for right), distance from the left telomere rounded to nearest kilobase, and the strand on which the SiRTA is located [(+) for the forward strand and (–) for the reverse strand].

To increase the throughput of this analysis pipeline and to map dnTA events with nucleotide precision, we developed the Pool-Tel-seq (PT-seq) method (*Materials and Methods* and Figure 1c). As in our original approach, a single inoculum is plated on a medium containing galactose to induce HO cleavage and thirty 5-FOA resistant colonies are identified that have lost the chromosome VII terminus. The 30 colonies are grown separately to saturation in liquid medium and equal volumes of each culture are pooled to generate a single genomic DNA sample for library construction and high through-put sequencing. The resulting sequence reads (>12 million) are analyzed for those that align at least partially to the 300 bp putative SiRTA and show evidence of telomere addition (TG_1-3_ or C_1-3_A sequence). To account for differences in read depth between experiments, the number of reads meeting these criteria is normalized to the number of reads mapping to a 300 bp sequence within *BRR6*, an essential gene on chromosome VII that lies centromere-proximal to the site at which the putative SiRTA is integrated (Figure 1c). At least two biological replicates are generated for each strain. To benchmark SiRTA efficiency based on our previous method, we applied PCR-based mapping and PT-seq to multiple 30-colony samples derived from SiRTAs of a range of efficiencies. Using linear regression, we find a strong correlation between the two methods (r^2^=0.97), allowing us to use this standard curve to estimate the number of colonies within a 30-colony sample that underwent dnTA at the putative SiRTA (Figure 1d, closed triangles; Supplementary File 3). This method also yields information about the relative frequency of telomere addition at each nucleotide position.

To verify that this method is applicable at other locations in the genome, we utilized PT-seq to test SiRTA 9L44(-) at its endogenous location on chromosome IX. *Cis*- and *trans*-acting mutations with effects on the efficiency of dnTA at SiRTA 9L44(-) were used to compare the PCR and PT-seq methodologies over a range of SiRTA efficiencies. Again, results of the two methods are strongly correlated (r^2^=0.95; Figure 1d, open circles; Supplementary File 3). The slopes of the standard curves generated at both chromosome locations are statistically indistinguishable (p = 0.76 by analysis of covariance), suggesting that the percent of GCR events incurring dnTA (SiRTA efficiency) can be accurately estimated from PT-seq results regardless of chromosome location.

### Putative SiRTAs are accurately identified using a computational method

Visual inspection of sequences found to function as SiRTAs revealed similarity to yeast telomeric sequences, consistent with prior work demonstrating that association of Cdc13 with the Stim sequence is required for dnTA (Obodo et al. 2016). The Core sequence is also TG-rich, likely reflecting required complementarity to the *TLC1* template sequence and (perhaps) the ability to associate with Cdc13. We postulated that SiRTA function could be predicted by considering not only the TG-richness of a sequence but also its similarity to the pattern of the yeast telomeric repeat (TG_1-3_). SiRTA function does not require a perfect match to the telomeric sequence, so we developed a strategy to score similarity to a telomeric repeat while allowing divergence from that pattern. The Computational Algorithm for Telomere Hotspot Identification (CATHI) identifies strings of consecutive Gs and Ts, awards one point for each base in that string, and subtracts 1.5 points for each instance of GGTGG, a sequence that is not found in yeast telomeres (Figure 2a). Calculations are done in a sliding window that can be varied in size (*Materials and Methods*).

**Figure 2.**
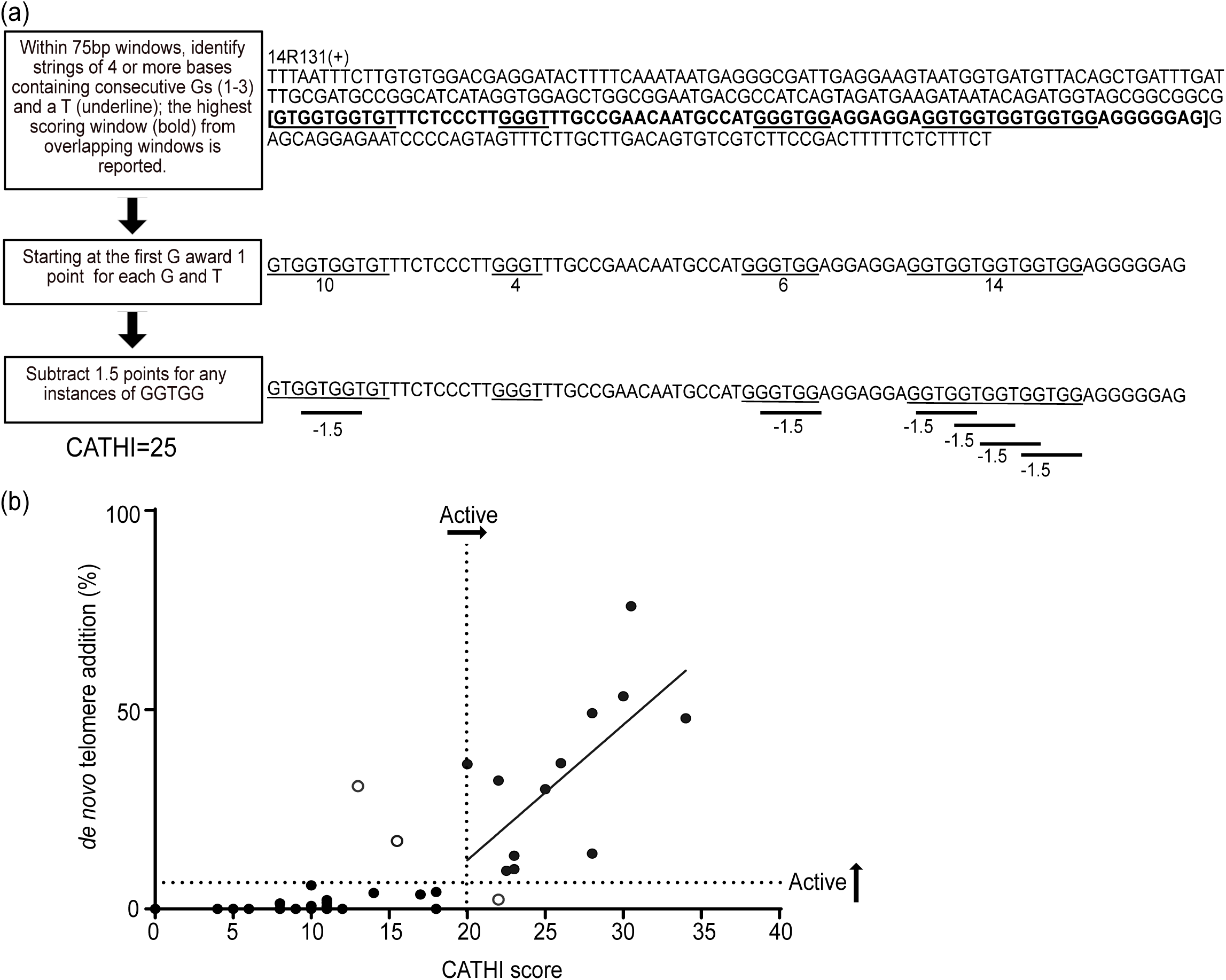
Computational Algorithm for Telomere Hotspot Identification (CATHI) predicts SiRTA function. a) Summary of methodology used to generate CATHI score. An example corresponding to SiRTA 14R131(+) is shown (300 bp sequence beginning at chromosome XIV nucleotide 131308). Although multiple 75 bp windows within this sequence surpass the threshold score of 20, the calculation is shown only for the highest-scoring window, starting at nucleotide 131471 (bold, bracketed text). Underlined sequences correspond to strings of 4 or more guanine or thymine nucleotides conforming to the patterns described in the flowchart and in more detail in *Materials and Methods*. Each underlined nucleotide is awarded one point. Subsequently, each occurrence of a GGTGG pentanucleotide incurs a 1.5-point penalty to generate the final score. b) Correlation of CATHI score and the percentage of GCR events that result from de novo telomere addition within the sequence of interest. Each value is the average of at least two experiments, each with 30 GCR events. The standard curve for chromosome VII (Figure 1d) is used to convert PT-seq values to the percentage of GCR events undergoing dnTA in the SiRTA. Horizontal dashed line indicates a minimum telomere-addition efficiency of 6.6% used to define an active SiRTA (see text for detail). Thirty-two of the sequences fall below this threshold and 15 are above this threshold. Vertical dashed line illustrates a CATHI score of 20 that effectively separates active and inactive sequences. Thirty-three sequences fall below this threshold and 14 are above this threshold (Supplementary File 3). The open circles are false negatives or false positives. Linear regression analysis on SiRTAs with a CATHI score of 20 or more yields a p-value of 0.01 (r^2^ =0.43).

The algorithm was developed through an iterative process in which sequences were identified and tested for SiRTA function. This dataset included several previously published SiRTAs, sequences identified during the work described here, and negative control sequences that were not expected to function as SiRTAs. To standardize measurements of SiRTA efficiency, all the tested sequences were assayed on chromosome VII by inserting a 300 bp region encompassing the putative SiRTA. If boundaries of the SiRTA sequence were previously established, the SiRTA was centered within the 300 bp region. All sequences were tested at least in duplicate and the average percent of clones undergoing telomere addition within the test sequence was determined in comparison to the chromosome VII standard curve (Figure 1d). During initial testing, data were obtained for 37 sequences, seven of which had an average SiRTA efficiency above the 6.6% cutoff. To optimize the algorithm, we calculated a score for each of the 37 sequences using varying window sizes (25 to 150 bp) and penalties (0 to 3) to identify a combination generating the best fit by linear regression (Supplementary File 4). A window size of 75 and a penalty of 1.5 for GGTGG sequences yielded the highest correlation between CATHI score and SiRTA efficiency (r^2^ = 0.63 for all sequences and 0.69 for the seven sequences exceeding the 6.6% cutoff for SiRTA function).

Testing of additional sequences after the algorithm parameters were established resulted in a final dataset of 47 sequences (13 active as SiRTAs) that are graphed relative to CATHI score in Figure 2b (Supplementary File 3). Using the threshold of 6.6% to define an active SiRTA, a CATHI score of 20 effectively separates active and inactive sequence with a false positive rate of ∼2% (1/47) and a false negative rate of ∼4% (2/47). We conclude that the algorithm can be used to accurately identify sequences with a propensity to stimulate dnTA. For those sequences with a CATHI score of 20 or greater, the score is moderately predictive of SiRTA efficiency (r^2^=0.43; p=0.015). The sequences with the four highest CATHI scores tested are also the most efficient. However, scores between 20 and 30 are less predictive of efficiency, suggesting that some aspects of SiRTA function are not captured by the algorithm (see *Discussion*).

### Distribution of SiRTAs across the yeast genome

Using the algorithm parameters established above, the 16 chromosomes (excluding any terminal TG_1-3_ telomeric sequences; see Supplementary File 2 for coordinates) were scanned as a series of 75 bp sliding windows with a step size of 1. Overlapping windows with scores of 20 or greater were merged such that the starting and ending coordinates of a predicted SiRTA represent the maximum distance between the first and last window meeting the threshold value. The final score assigned to a set of overlapping windows is equivalent to the highest CATHI score in that set. The algorithm was separately applied to the top and bottom strands and strand information was retained. Overall, we identified 728 sequences in the *S. cerevisiae* genome with a CATHI score of 20 or greater (Supplementary File 4).

We examined the overall distribution of these 728 sequences within the 16 yeast chromosomes (Figure 3). SiRTAs on the top strand (5’ to 3’; 342) are shown in blue and those on the bottom (3’ to 5’; 386) strand are shown in red (Figure 3b and c). The centromere of each chromosome is depicted as a black circle. The overall distribution between the two strands is not different from random (p=0.25 by chi-square test). However, there is a minor but significant tendency for the SiRTAs to cluster on the same strand. This effect is quantified by measuring the number of times a SiRTA is found on the opposite strand from its neighbor (number of “strand switches”). We observe 322 strand switches across the genome, significantly fewer than the number expected by chance (351.8 ± 13.0; p=0.013; Supplementary Figure 2a). This difference is driven almost entirely by a reduction in the number of “singlet” SiRTAs compared to what would be expected by chance (148 observed versus 187.7 ± 15.0 expected; p=0.0043; Supplementary Figure 2b). In contrast, runs of longer length do not deviate significantly from expectation. We conclude that there is a minor tendency for SiRTAs to cluster on the same strand, but only with the nearest neighbor.

**Figure 3.**
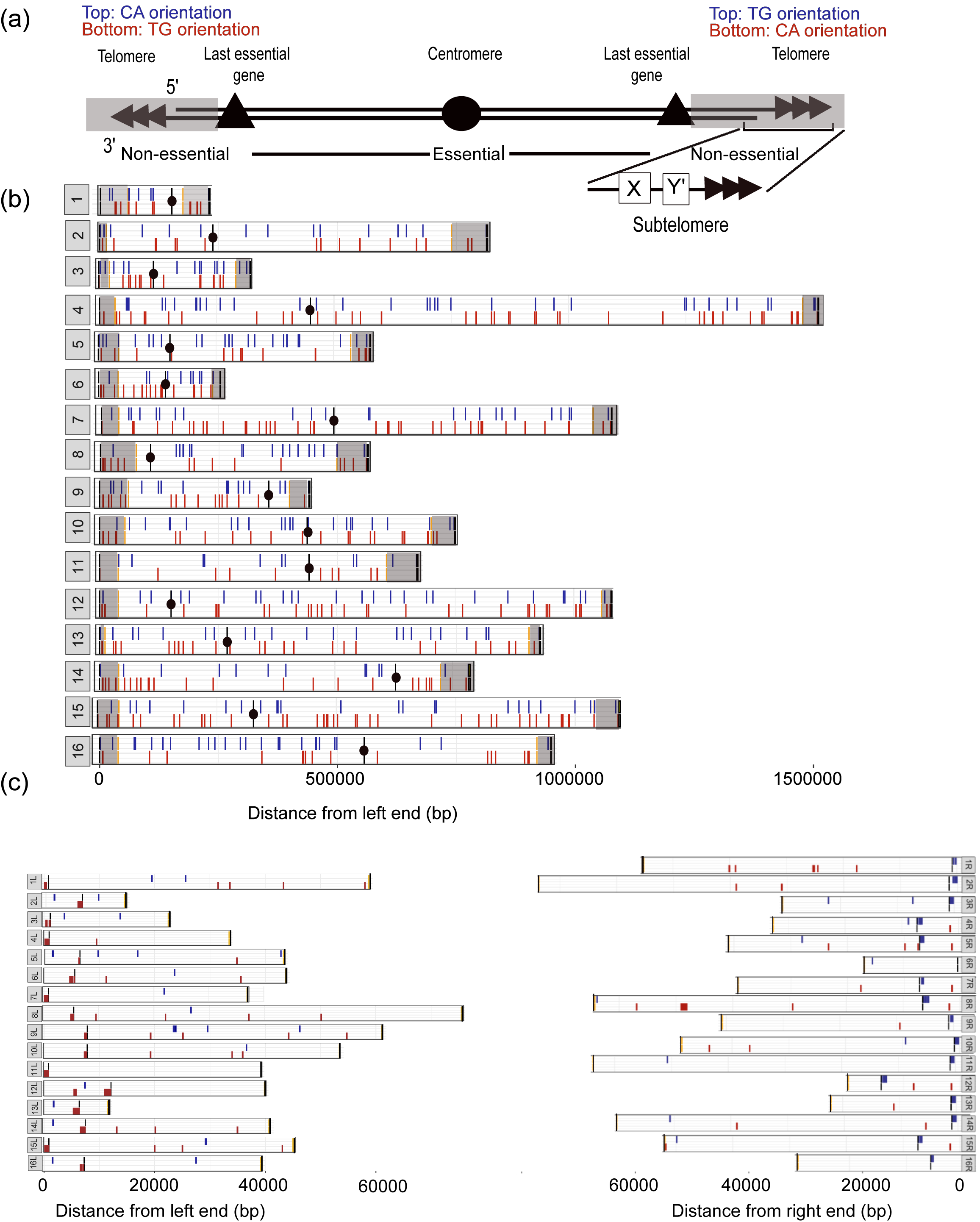
Summary of predicted SiRTAs across the *S. cerevisiae* genome. a) Diagram of chromosome landmarks. Predicted SiRTAs are listed in Supplementary File 4. Non-essential regions (grey boxes) are sequences located between the last essential gene (most distal gene on the chromosome arm that causes lethality in a haploid strain when deleted) and the telomere on each chromosome arm. Subtelomeres are located immediately adjacent to the telomeric repeats within the nonessential regions and are composed of a single complete or partial X element (all telomeres) and one or more Y’ elements (a fraction of telomeres). Genomic coordinates are listed in Supplementary File 2. Location of the CA- or TG-oriented sequences are indicated for the left and right chromosome arms (see text). b) Distribution of SiRTAs on each of the 16 yeast chromosomes; distance in base pairs from the left telomere is indicated at the bottom of the figure and corresponds to coordinates in the S288C reference genome. Black circles mark the centromere position and nonessential regions are highlighted in grey. Blue lines in the top half of each bar represent SiRTAs on the top (plus) strand and red lines in the bottom half of each bar refer to SiRTAs on the bottom (minus) strand. c) Distribution of SiRTAs in the nonessential regions of the left and right arms of each chromosome as defined in Supplementary File 2. The transition between the subtelomeric X element and the remainder of the nonessential region is shown as a horizontal black line. Red and blue lines are as described in (b). Diagrams in (a) and (b) were generated using shinyChromosome (Yu et al. 2019).

In a haploid cell, chromosome truncation proximal to the last essential gene on a chromosome arm will be lethal. Therefore, we were interested in determining whether the distribution of putative SiRTAs is different in essential versus nonessential regions. For our analysis, we defined the nonessential region on each chromosome arm as comprising sequences distal to the last essential gene (Figure 3a; Supplementary File 2). The last essential gene, in turn, is the most telomere-proximal gene for which single gene deletion was reported to cause lethality during the systematic knockout of each open reading frame in *S. cerevisiae* (Giaever et al. 2002). This definition does not take synthetic lethality into account; some nonessential regions may be smaller than defined here if the combined loss of one or more genes in that region results in cell death. In Figure 3b, nonessential regions are highlighted in gray; those same regions are shown in expanded form in Figure 3c. The nonessential regions are divided into sequences that are unique (in most cases) among the different chromosome arms and the highly repetitive subtelomeric X and Y’ elements found immediately adjacent to the telomeric repeats (Figure 3a). All chromosome arms contain at least a portion of the X element while some also contain one or more Y’ elements (∼6 kb each; Louis and Haber 1990; Zhu and Gustafsson 2009). The transition to subtelomeric sequence is shown on each chromosome arm as a black line (Figure 3c). Diagrams in Figure 3 are based on the published sequence of reference strain S288C. Recent long-read sequencing analyses confirm that some subtelomeric regions contain additional terminal sequences (primarily Y’ elements) that were not included in the published reference genome (Yue et al. 2017), so our analysis likely underestimates the number of subtelomeric SiRTAs.

To test the hypothesis that SiRTAs are preferentially located in nonessential terminal regions, we utilized a permutation-based enrichment test to compare the distribution of SiRTAs between essential and non-essential regions to that of randomly shuffled sequences matched in number and length to the sequences identified by the CATHI algorithm. This analysis shows significant enrichment for SiRTAs in the nonessential regions of the genome (p<0.01) but this enrichment disappears when the subtelomeric regions (X and Y’ elements) are excluded from the analysis (Figure 4a). We conclude that SiRTAs are disproportionately enriched in nonessential regions, but that this effect is limited to the most distal, highly repetitive subtelomeric sequences. Seventy-five of the 728 putative SiRTAs lie within the subtelomeric sequences. Nine of those 75 sequences consist of perfect telomeric (TG_1-3_) repeats located between X and Y’ elements and seven of these perfect repeats constitute the top scoring sites in the genome (Figure 4b and Supplementary File 5). The remaining predicted SiRTAs in subtelomeric regions are not comprised of perfect telomeric repeats and lie within the X or Y’ elements. We were concerned that the tracts of perfect telomeric repeats might affect our analysis. However, when the nine perfect telomeric repeats are excluded, there remains significant enrichment for SiRTAs within the nonessential regions (p=0.001; Supplementary Figure 3). Notably, all chromosome ends [with one exception: 6R)] contain at least one region predicted to function as a SiRTA (Figure 3c). We conclude that SiRTAs are disproportionately found within the subtelomeric regions but are otherwise not significantly enriched within nonessential sequences.

**Figure 4.**
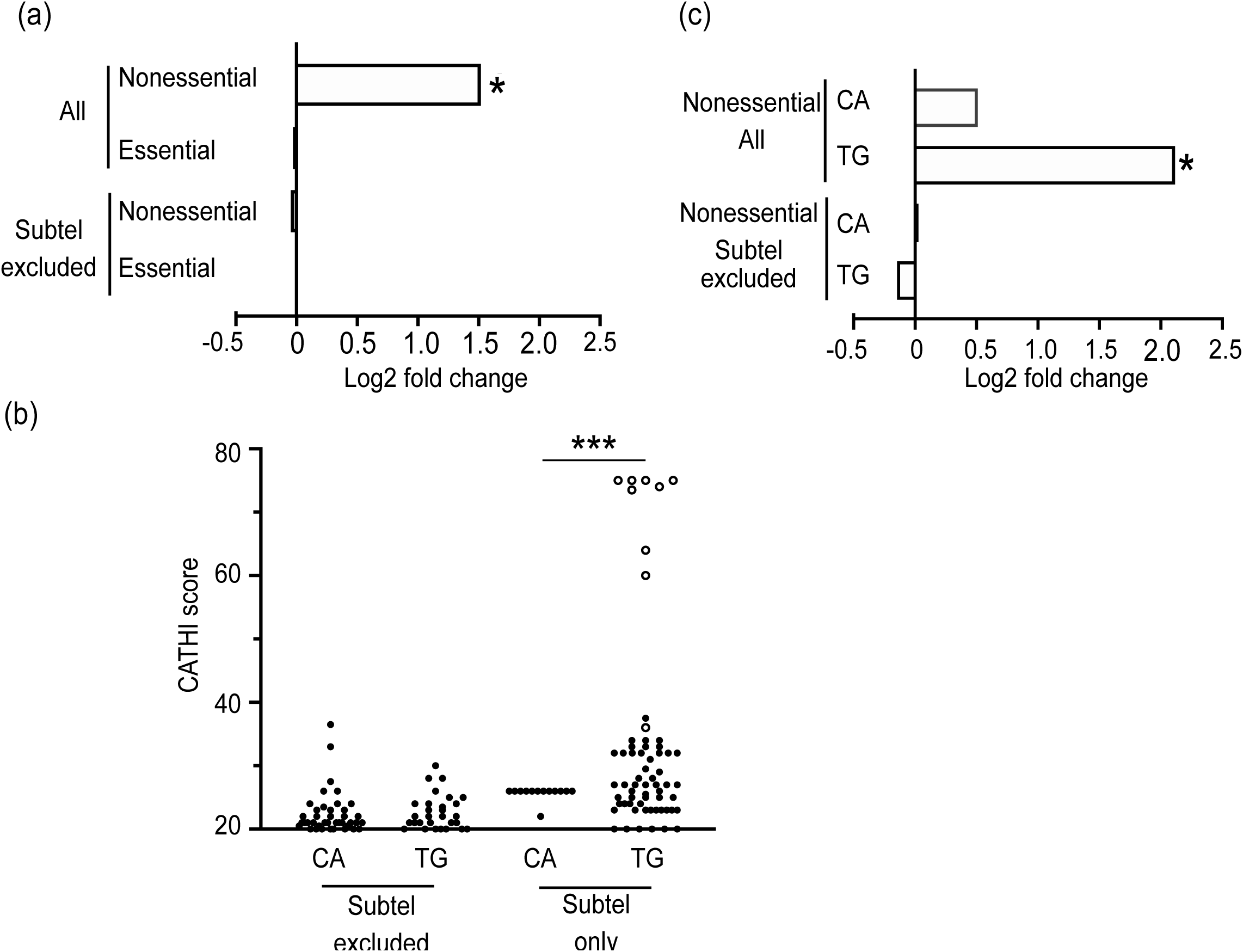
SiRTAs are enriched in subtelomeric regions. a) Using a permutation strategy as described in *Materials and Methods*, the enrichment of SiRTAs (Log2 fold change) was determined for the nonessential and essential chromosome regions. Analysis utilized all genomic sequences (except the terminal telomeric repeats) or genomic sequences from which the subtelomeric regions were excluded, as indicated (coordinates in Supplementary File 2). *p-value <0.01 by chi-squared test with Bonferroni’s correction.  b) Distributions of CATHI scores (20 and greater) for putative SiRTAs in the TG- or CA-orientations. Analysis is presented separately for SiRTAs in nonessential regions (subtel excluded) versus those in the subtelomeric repeats (subtel only). Nine SiRTAs containing perfect telomeric (TG_1-3_) repeats are indicated with open circles. ****p <0.0001 by chi-squared test. Coordinates of telomeric repeats are found in Supplementary File 5.  c) Enrichment analysis was conducted separately for SiRTAs in the TG or CA orientation (see text and Figure 3a for definitions) as described in part (a). Results for the nonessential regions are shown. *p-value<0.01 by chi-squared test with Bonferroni’s correction.

As described in the *Introduction*, the strand on which a SiRTA is located has important implications for the consequence of dnTA. SiRTAs in the TG-orientation (those oriented to stabilize the centromere-containing fragment when a break occurs distal to the site) are on the bottom strand for the left arm of a chromosome and on the top strand for the right arm. SiRTAs in the genome as a whole do not show a bias for the TG- versus CA-orientation (372 versus 356; p=0.675 by chi-square test). To examine the distribution more carefully, we examined SiRTA orientation within the nonessential regions, where an excess of SiRTAs was already noted (Figure 4a). Interestingly, enrichment within nonessential regions is observed only for SiRTAs in the TG-orientation (p<0.01), while SiRTAs in the CA-orientation are not significantly enriched (Figure 4c). The same result is observed when the nine perfect TG_1-3_ repeats are removed from the analysis (p=0.001; Supplementary Figure 3). As expected, enrichment is no longer observed when subtelomeric sequences are excluded (Figure 4c). This differential distribution is visualized in a plot showing the CATHI score of predicted SiRTAs within the nonessential chromosome regions. In non-subtelomeric regions, distributions are indistinguishable for the two orientations (Figure 4b). In contrast, SiRTAs in the X and Y’ elements are much more likely to be in the TG-orientation than in the CA-orientation (p<0.0001 by chi-square test; Figure 4b). The enrichment of TG-oriented SiRTAs within subtelomeres is also visually apparent in the clustering of red symbols at or near the subtelomere junction on all left arms and of blue symbols on nearly all right arms (except 6R; Figure 3c).

We tested the ability of three sequences identified in subtelomeric regions to stimulate dnTA using the HO cleavage assay. Two of these sequences overlap with an X element [14L07(-) and 15R1084(+)] and one overlaps with a Y’ element [7R1089(-)]. All three sequences function as SiRTAs, stimulating de novo telomere addition at frequencies of 10.0%, 13.3% and 36.6%, respectively (Supplementary Figure 4). Although 7R1089(-) functions well as a SiRTA in our assay, it is worth noting that it is in the CA orientation. Because it is part of a conserved Y’ sequence, a very similar sequence occurs at multiple chromosome ends (also in the CA orientation). The X element sequences are TG-oriented and represent sequences found in two distinct regions of the X element sequence. Taken together, these results support the interesting possibility that sequences capable of functioning as SiRTAs have been retained near chromosome termini to facilitate chromosome rescue in the event of telomere loss.

### TG-rich sequences identified by the algorithm are overrepresented in the yeast genome

We were curious whether sequences predicted to stimulate dnTA are found in yeast at the expected frequency given the nucleotide composition of the yeast genome. To address this question, we generated five scrambled genomes identical in sequence composition to the yeast genome. To avoid potential biases introduced by the subtelomeric regions, this analysis was done on sequences from which the subtelomeric X and Y’ elements were excluded (see *Materials and Methods*). The scrambled genomes contain, on average, 283.2 ± 18.3 sequences that score 20 or higher compared to 653 sequences observed in the yeast genome, an excess of 2.3-fold. The differential becomes increasingly apparent at higher scores with an excess of 1.8-fold at a score of 20 and an excess of 5.2-fold at a score of 25. Among the scrambled genomes, an average of fewer than one sequence has a score of 30 or higher (range 0 to 2), while 21 sequences scoring 30 or higher are observed in the *S. cerevisiae* genome (Figure 5a and Supplementary File 6). In Figure 5b, putative SiRTAs with scores of 27 or higher are shown to emphasize the strikingly different distributions in the simulated versus actual genomes. To address whether the excess of higher scores might be functionally related to SiRTA function, we examined the predicted and actual occurrence of CATHI scores less than 20, which are unlikely to stimulate increased levels of dnTA (see Figure 2b). For sequences with CATHI scores of 15-19, we still observe an excess in the actual genome, although the excess is less pronounced (1.3-fold, with 2703 ± 16.3 predicted compared to 3485 observed; Figure 5c).

**Figure 5.**
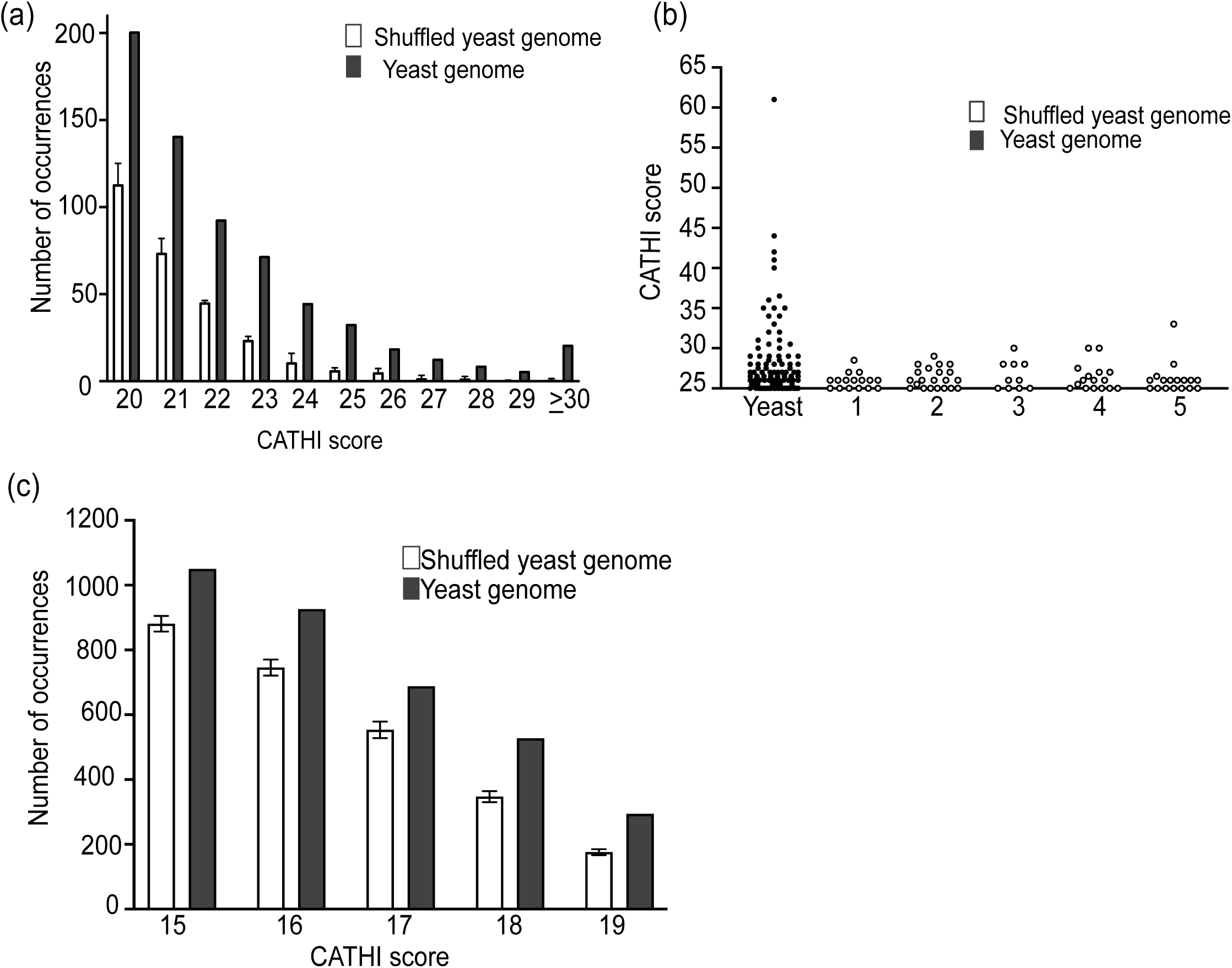
CATHI scores are significantly elevated in the *S. cerevisiae* genome relative to expectation. a) As described in *Materials and Methods*, the algorithm was applied to the *S. cerevisiae* genome (excluding subtelomeric regions) and the number of sequences at each score (15 or higher; rounded down to the nearest integer) was graphed (solid bars). Genomic sequences (excluding subtelomeric regions) were scrambled five times and the identical procedure was applied (Supplementary File 6). Data are presented as the average and standard deviation of the five trials (open bars). b)  Distribution of CATHI scores of 25 and above in the *S. cerevisiae* genome (closed circles) and shuffled genomes (open circles). Subtelomeric sequences were excluded. c) As in (b), but data are shown for CATHI scores ranging from 15-19.

### TG-dinucleotide repeats stimulate dnTA and are among the strongest SiRTAs in the genome

In analyzing predicted sites of dnTA, our attention was particularly drawn to SiRTA 6R210(+). With a score of 61, this sequence represents the strongest predicted site outside the subtelomeric regions (see outlier in Figure 5b). SiRTA 6R210(+) contains a nearly perfect 62 nucleotide TG-dinucleotide repeat and is the longest TG-dinucleotide repeat in the *S. cerevisiae* genome (the next longest is 41 nt; Supplementary File 5). Interestingly, the sequence is in the TG-orientation but lies centromere-proximal to the last essential gene on the left arm of chromosome VI, implying that repair by dnTA at this site in a haploid cell would be lethal.

To determine the efficiency of dnTA at SiRTA 6R210(+), we inserted the TG-dinucleotide repeat (centered within a 300 bp region) at the test site on chromosome VII. Most strains that we monitor for SiRTA function on chromosome VII generate GCR events at a frequency of ∼0.001%, equivalent to a negative control strain lacking a SiRTA. In contrast, a strain containing the 62 bp TG-dinucleotide repeat generates 5-FOA resistant colonies at a 10-fold higher frequency of ∼0.01% (Figure 6a). By PT-seq, 86% of GCR events involve dnTA addition within the inserted sequence (Figure 6b). Therefore, although the percent of events at the SiRTA that involve dnTA is similar in strains carrying the TG-dinucleotide repeat versus the efficient SiRTA 14L35(-) (76%), the overall frequency with which dnTA occurs at the 62-nt dinucleotide repeat is at least 10 times higher. We conclude that SiRTA efficiency alone (defined as the percentage of GCR events in which telomere addition occurred within the sequence of interest) underestimates the propensity of a sequence to stimulate telomere addition when SiRTA activity is very high. For this reason, we did not include SiRTA 6R210(+) in the comparison of CATHI score and SiRTA efficiency in Figure 2b.

**Figure 6.**
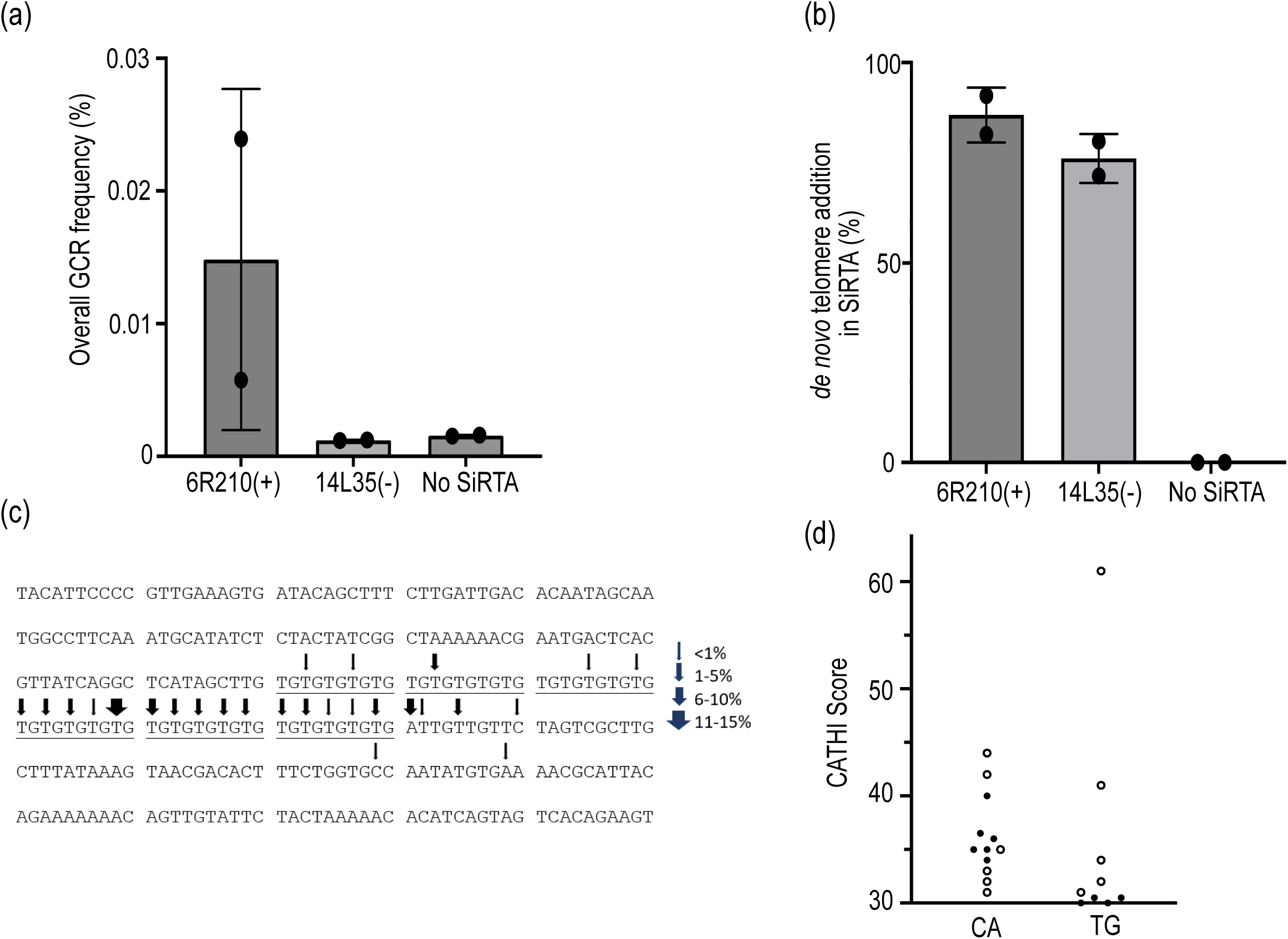
A 62 bp TG-dinucleotide repeat [SiRTA 6R210(+)] supports high levels of dnTA. a) Strains in which a 300 bp sequence encompassing SiRTA 6R210(+) or 14L35(-) was integrated on chromosome VII were subjected to the HO-cleavage assay as described in *Materials and Methods*. The percent of cells that survived on galactose-containing medium and acquired 5-FOA resistance [cells containing a gross chromosomal rearrangement (GCR)] is shown. A strain lacking any insertion (No SiRTA) was utilized as a control. Error bars are standard deviation. b) The percent of GCR events that involve de novo telomere addition in the sequence of interest was determined by PT-seq for each strain described in (a). Each data point was generated by analysis of 30 GCR events. Average and standard deviation are shown. c) The 300 bp sequence encompassing 6R210(+) is shown. Sequence reads generated by PT-seq from a total of 60 GCR events [corresponding to the experiments shown in (b)] were filtered for those containing evidence of de novo telomere addition within the 300 bp sequence. Sites at which de novo telomere addition was observed are indicated (arrows). Arrow width indicates the percent of telomere-containing reads that map to that particular site.  d) CATHI scores are shown for all non-subtelomeric SiRTAs with scores of 30 or higher, separated by CA- or TG-orientation. Open circles correspond to SiRTAs containing TG-dinucleotide repeats (also listed in Supplementary File 5).

Using the PT-seq results, we mapped the sites at which dnTA occurred at SiRTA 6R210(+) (Figure 6c). Each arrow corresponds to the last nucleotide that aligns between the chromosome and at least one PT-seq read, representing the 3’-most nucleotide at which synthesis by telomerase may have initiated. Given that ∼86% of the 60 individual strains represented in the analysis contain evidence of dnTA, these results correspond to the mapping of 50-52 independent telomere addition events. Sites identified in a larger fraction of reads were likely targeted by telomerase in multiple independent clones. Consistent with our prior observation that a 5’ Stim sequence is required to stimulate telomere addition in a 3’ Core sequence (polarity relative to the TG-rich strand), the vast majority of telomere addition events occur in the 3’ half of the dinucleotide repeat or in sequences immediately downstream. Two events occurred at least 50 bases downstream of the TG-repeat, consistent with prior reports that TG-rich sequences can stimulate dnTA within neighboring sequences (Kramer and Haber 1993; Mangahas et al. 2001).

Excluding the subtelomeric regions, TG-dinucleotide repeats comprise 11 of the 21 CATHI scores of 30 or greater (Figure 6d). SiRTA 7L69(-) (CATHI score = 34) contains a 34-nt perfect TG repeat and was identified by the Zakian laboratory as a site capable of stimulating dnTA in response to a DSB induced more than 50kb distal to the eventual site of telomere addition (Mangahas et al. 2001). When integrated at the test site on chromosome VII, this 34-nt repeat stimulates dnTA with an efficiency of 47.9% by PT-seq (Supplementary File 3). Together, these observations focus attention on TG-dinucleotide repeats as potential mediators of genome instability.

### Sequences that function to stimulate de novo telomere addition bind Cdc13 in vitro

Previous studies demonstrated that Cdc13 binding at the Stim sequence is required to promote dnTA. We hypothesized that sequences with CATHI scores of 20 or greater will bind Cdc13 with greater affinity than sequences with lower scores. Additionally, we predicted that the two sequences identified as false negatives in our initial analysis (Figure 2b) should bind Cdc13 with higher affinity than the single sequence identified as a false positive. To test these predictions, we utilized fluorescence polarization to measure the ability of unlabeled, 75-base oligonucleotides to reduce association of the Cdc13 DNA binding domain (Cdc13-DBD) with a 6-carboxyfluorescein (FAM) labeled 11-mer containing the canonical Cdc13 binding site (5’-GTGTGGGTGTG; referred to here as Tel11). Cdc13-DBD binds with similar sequence specificity and affinity to the Tel11 sequence as full-length Cdc13 (Lewis et al. 2014). The goal of these analyses was not to identify individual Cdc13 binding sites, but rather to measure the relative, cumulative ability of each sequence to bind Cdc13.

We first established that the FAM-labeled Tel11 oligonucleotide binds Cdc13-DBD (Supplementary Figure 5a) and selected concentrations of 30 nM Cdc13-DBD and 25 nM labeled Tel11 for the competition analyses. The apparent inhibition constant (*K*_i,app_) is defined as the concentration of each competitor required to reduce binding to the FAM-labeled Tel11 by half. An unlabeled 75-mer containing the Tel11 sequence at the center of the oligonucleotide (Tel11-75) was included in each experiment and normalized *K*_i,app_ values are reported as fold change relative to this control (*K*_i,app_ of Tel11-75/K_i,app_ of experimental oligonucleotide). Sequences flanking the Cdc13 consensus binding site in Tel11-75 lack any TG or GG motifs to minimize additional association of Cdc13-DBD. Oligonucleotides used in these assays are found in Supplementary File 1 and representative competition curves are shown in Supplementary Figure 5b. To validate the method, we determined the normalized *K*_i,app_ of a double-stranded version of Tel11-75 (0.5 +/−0.3) and the inverse complement of the Tel11-75 sequence (0.4 +/−0.2), both of which show the expected reduction in binding relative to Tel11-75 (Figure 7a). A 75-mer sequence from chromosome VI previously shown to lack SiRTA activity (CATHI score=5) also competes very weakly for Cdc13-DBD association (normalized K_i,app_ = 0.5 +/−0.2; Figure 7a). Finally, as expected, a 2xTel11-75 sequence that contains two adjacent Tel11 sequences competes twice as well as the Tel11-75 control oligonucleotide (normalized K_i,app_ = 2.2 +/−0.2; Supplementary Figure 6a).

**Figure 7.**
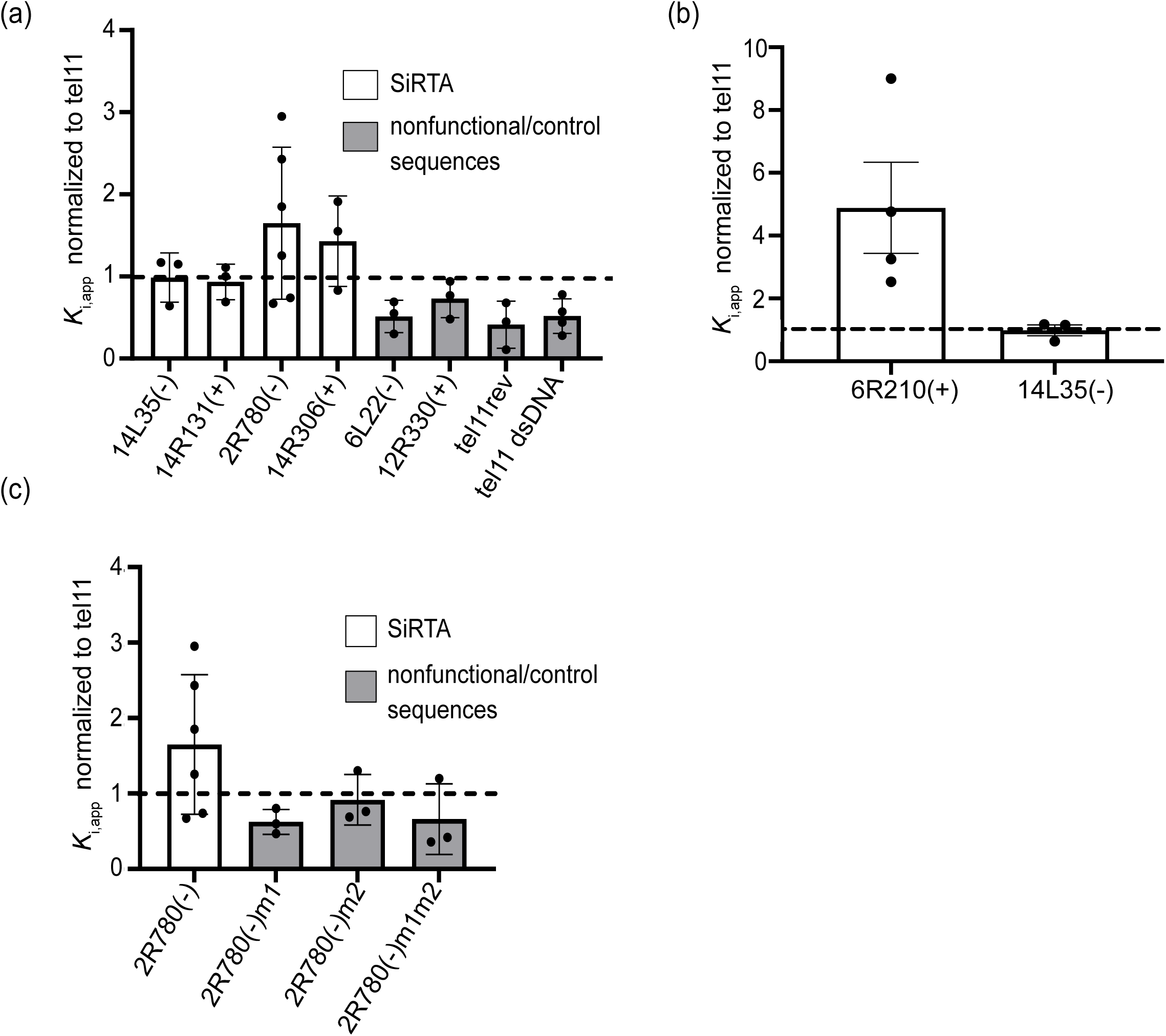
Sequences that function as SiRTAs bind Cdc13 *in vitro.* a) A competition fluorescence polarization assay was utilized to measure the relative association of the Cdc13 DNA binding domain with the indicated sequences (Supplementary File 1). Relative *K*_i,app_ was determined as described in *Materials and Methods*. Each point represents an independent measurement; error bars are standard deviation (Supplementary File 7). The dotted line indicates normalization of values to the *K*_i,app_ of a tel11-75 oligonucleotide included in each experiment. b) Same as in (a). *K*_i,app_ of the TG-dinucleotide repeat analyzed in Figure 6 is shown [6R210(+)]. Data for 14L35(-) are repeated from (a) for comparison. c) Same as in (a). 2R780(-) and its mutated variants are described in Hoerr et al. (2023).

We chose to test several sequences with CATHI scores over 20 that had been previously shown to stimulate dnTA. Oligonucleotides were designed to correspond to the 75 bases with the highest CATHI score within the 300bp sequence tested for SiRTA function. Both 14L35(-) and 14R131(+) compete in a manner indistinguishable from the Tel11-75 control sequence (normalized *K*_i,app_ of 1.0 +/−0.3 and 0.9 +/−0.2, respectively) and bind more robustly than the negative control sequences (Figure 7a). The two sequences identified as false negatives in Figure 2b [2R780(-) and 14R306(+)] both compete more effectively than the Tel11-75 control sequence (normalized *K*_i,app_ of 1.6 +/−0.9 and 1.4 +/−0.6, respectively; Figure 7a). This observation is consistent with the ability of these sequences to stimulate dnTA and suggests that the algorithm fails to predict Cdc13 binding in some cases. We also tested the false positive sequence [12R330(+)] with a CATHI score of 22 and an average dnTA frequency of 2.3% (below our cut-off for SiRTA function). This 75-base sequence has a normalized *K*_i,app_ of 0.7 +/−0.2, intermediate to that of the Tel11-75 control sequence and the negative controls (Figure 7a).

Given the extremely high SiRTA activity of the 62-nt TG-dinucleotide repeat described above, we tested the ability of a 75-mer containing this repeat to compete for Cdc13-DBD binding. The normalized *K*_i,app_ of 4.9 +/−2.9 measured for this sequence is considerably higher than any other sequence tested (Figure 7b). The first, second, and fourth base of the canonical Cdc13 binding site (5’-**<u>G</u>**T**<u>GT</u>**GGGTGTG) contribute most strongly to Cdc13 affinity, comprising a GxGT motif that recurs in the TG-dinucleotide motif. The 62-nt dinucleotide repeat is predicted to accommodate approximately five 11-mer binding sites, remarkably close to the observed 4.9-fold increase in competition compared to the Tel11-75 control oligonucleotide with a single binding site.

Our prior analysis of SiRTA 2R780(-) presented an additional opportunity to test the correlation between Cdc13 binding and SiRTA efficiency (Hoerr et al. 2023). SiRTA 2R780(-) contains a Stim sequence of ∼50 bases, deletion of which abrogates dnTA. We previously showed that mutation of either one of two GxGT motifs located in this 50 nt Stim sequence greatly diminishes SiRTA function, an effect that we attributed to reduced Cdc13 association (Hoerr et al. 2023). Consistent with this hypothesis, we find that mutation of one or both motifs significantly reduces the ability of the oligonucleotide to compete for Cdc13-DBD binding (Figure 7c).

The experiments described above provide evidence that sequences capable of stimulating dnTA associate more robustly with Cdc13 than sequences that do not function as SiRTAs, although our ability to distinguish borderline cases is limited. While there appears to be a threshold of binding required for SiRTA function, the cumulative “affinity” of a sequence measured in this assay is not fully predictive of SiRTA efficiency [e.g. 14L35(-) and 14R131(+) compete equivalently, but differ by a factor of two in SiRTA efficiency; 80.5% versus 30.7%]. This discrepancy may result in part from our choice of 75-mer sequence to test in each case, but also likely reflects the specific number, affinity, and distribution of Cdc13 binding sites within the sequence.

As another approach to benchmark the effect of high affinity Cdc13 binding on SiRTA function, we tested the ability of either a single canonical Cdc13 binding site or two tandem sites to stimulate dnTA at the test site on chromosome VII. One copy of the Tel11 site stimulated telomere addition in only four or five of 60 GCR events analyzed by PT-seq (7.5%; Supplementary Figure 6b). Remarkably, adding a second Tel11 (2xTel11) site increases the frequency GCR events undergoing dnTA to 83.3% (Supplementary Figure 6b). Together these results demonstrate the importance of Cdc13 binding sites for stimulating dnTA at SiRTAs.

### SiRTA distribution is not strongly associated with known protein binding sites or chromosome landmarks

To gain insight into factors that may contribute to dnTA, we examined whether putative SiRTAs preferentially overlap with the binding sites of proteins related to telomere addition such as Est2 (a component of telomerase recently shown to associate with internal chromosome sites; Lendvay et al. 1996; Pandey et al. 2021), Rap1 (a transcription regulator that also binds telomeric repeats and affects telomere length homeostasis; Conrad et al. 1990; Rhee & Pugh, 2011), and Pif1 (a helicase that negatively regulates telomerase at telomeres and DNA double-strand breaks; Schulz and Zakian 1994; Paeschke et al. 2011). We also examined the correlation between predicted SiRTAs and fragile sites, identified as regions that associate with γH2AX even in the absence of exogenous damage (Downs et al. 2000; Capra et al. 2010), and between SiRTAs and sequences predicted to form G-quadruplex structures (Capra et al. 2010). Using a permutation-based enrichment test under conditions that require overlap with at least half of the predicted SiRTA sequence, we find statistically significant enrichment among SiRTAs for Est2, Rap1, γH2AX binding sites, and G4-forming sequences (Supplementary Figure 7a and b). However, overlap never exceeds 20% of the putative SiRTAs, arguing against a strong functional relationship. Although the actual number of putative SiRTAs overlapping with a Pif1 binding site is the highest of all characteristics tested (18.4%), this extent of overlap is not significant, likely because the regions reported to bind Pif1 by chromatin immunoprecipitation are relatively broad. To determine whether the predicted strength of the SiRTA affects these results, we divided putative SiRTAs into tertiles based on CATHI score but found no strong relationship between CATHI score and the significance of overlap (Supplementary Figure 7c). There is also no obvious enrichment among sequences that are known to function as SiRTAs. Of the 14 SiRTAs known to be active, only one overlaps with an Est2 binding site and one overlaps with a Rap1 binding site. Overall, these results fail to identify any overlapping binding sites with evidence of strong functional significance and suggest that fragile sites and G-quadruplex forming sequences are not strongly correlated with predicted hotspots of dnTA.

### SiRTAs are predominantly found within coding regions

We were interested in determining whether SiRTAs are preferentially excluded from genic regions due to higher levels of evolutionary constraint. Each of the 728 SiRTAs was categorized as genic (any part of the SiRTA overlapped with a gene as defined by the start and stop codon of each annotated gene) or intergenic (Supplementary File 9). Seventy-eight percent of all SiRTAs overlap with coding regions and only 22% are exclusively found in intergenic regions (Supplementary Figure 8). Given that approximately 30% of the yeast genome is intergenic (Hurowitz and Brown 2003; Lynch et al. 2008), we conclude that SiRTAs are not excluded from expressed regions. There are two exceptions. When a SiRTA contains long (>20 nt) TG-dinucleotide repeats, those repeats are virtually never found within a coding region. This result is not surprising since expansion or contraction of a dinucleotide repeat is expected to disrupt the open reading frame. Second, intergenic SiRTAs are disproportionately found in the subtelomeric X and Y’ elements. While only 8% of all SiRTAs are in the subtelomeric regions, 37% of intergenic SiRTAs are subtelomeric, consistent with the presence of few transcribed regions in the subtelomeres (Supplementary File 9). Interestingly, for those SiRTAs that overlap with open reading frames, 58% are located on the template strand, which is different from the expectation of random distribution (Supplementary Figure 8; p<0.05 by Chi square test) and suggests that the presence of these sequences within genes may, in some cases, have consequences for cellular fitness.

## Discussion

### Prediction of SiRTA function

In this work, we predict the distribution of SiRTAs in the yeast genome, an important step in understanding the role of these sequences in genome stability and function. The Zakian laboratory initially proposed that hotspots of dnTA addition contain tracts of 15 or more nucleotides consisting exclusively of T and G in a “telomere-like” pattern (Mangahas et al. 2001). Our subsequent analysis of SiRTAs on chromosome V and IX revealed that these requirements are too strict. For example, SiRTA 5L35(-) (formerly called 5L-35) stimulates dnTA in response to both spontaneous and induced DSBs (Stellwagen et al. 2003; Obodo et al. 2016; Ngo et al. 2020), but the longest uninterrupted string of TG sequence is 14 nucleotides, including several instances of a TT motif that never occurs within telomeric repeats.

Given this information, we set out to develop a method that could reliably predict whether a particular sequence has the capacity to stimulate unusual levels of dnTA (Figure 2). The algorithm described here prioritizes “telomere-like” sequences, but provides flexibility for some deviation from that pattern. With a single exception (discussed below), sites previously identified to stimulate dnTA are predicted by the algorithm to function as SiRTAs. For example, the Zakian lab identified three sites on chromosome VII that stimulate dnTA following an induced DSB (Mangahas et al. 2001). Two of these sites, now renamed SiRTA 7L67(-) and 7L69(-) (CATHI scores of 28 and 34, respectively), were originally found to stimulate dnTA at a distance of more than 50 kb from the induced DSB. In the standardized conditions of our chromosome VII test site where the break is induced ∼2 kb from the sequence of interest, these SiRTAs stimulate dnTA with efficiencies of 49.1% and 47.9% (Figure 2b and Supplementary File 3). The third site identified by the Zakian lab lies within the *URA3* gene, integrated at an ectopic location internal to the induced break. Although we did not test this sequence in our assay, it has a CATHI score of 22 and is annotated as SiRTA 5R117(+) to reflect the native location of *URA3* on chromosome V (Supplementary File 4).

Overall, the algorithm presented here correctly predicts SiRTA function (yes or no) with an accuracy close to 95% (44/47). Because false positives and false negatives occur at similar frequency, our estimate of ∼650 SiRTAs (excluding subtelomeric X and Y’ elements) is likely quite accurate, based on the definition of a SiRTA proposed here.

### Sequences that stimulate dnTA associate with Cdc13

Because prior work suggests that dnTA is stimulated by association of Cdc13 with single-stranded DNA generated after a DSB, the CATHI algorithm likely identifies sequences with affinity for Cdc13. To test this hypothesis, we developed a fluorescence polarization competition assay in which sequences are tested for their relative ability to compete with a labeled oligonucleotide for binding to the purified Cdc13 DNA binding domain. A 75-mer oligonucleotide containing two tandem copies of the canonical Cdc13 binding site competes twice as well as an oligonucleotide containing a single site, suggesting that the assay is sensitive to Cdc13 binding (Supplementary Figure 6a). Our goal is to measure overall association of Cdc13, which arises as a combination of the number and affinity of binding sites. We find that 75-mer oligonucleotides containing sequences that stimulate dnTA compete as well or better than a 75-mer containing a single match to the telomeric consensus Cdc13 binding sequence. In contrast, sequences that fail to support dnTA compete less well (Figure 7a). Importantly, mutations previously demonstrated to reduce SiRTA function also reduce Cdc13 binding (Figure 7c). Together with our previous demonstration that dnTA is stimulated through the artificial recruitment of Cdc13 to the Stim sequence of a SiRTA (Epum et al. 2020; Hoerr et al. 2023; Obodo et al. 2016), these results are consistent with the requirement for a threshold level of Cdc13 in stimulating dnTA.

Despite the observation at a single SiRTA that Cdc13 association correlates well with SiRTA efficiency, the apparent overall affinity for Cdc13 measured by fluorescence polarization poorly predicts SiRTA efficiency. For example, SiRTA 14L35(-) competes equivalently with the control sequence but stimulates dnTA more strongly than most other SiRTAs, including those that compete more effectively for Cdc13 binding. This apparent discrepancy may reflect an effect of the distribution or spacing of Cdc13 binding sites on SiRTA function. In prior work, we observed that deletion of the ∼30 nt spacer region between the Stim and Core sequences of a SiRTA dramatically increases SiRTA efficiency (Obodo et al. 2016). The highly efficient SiRTA 14L35(-) contains an unusually long region of TG-rich sequence that likely acts as both a Stim and Core region with little or no spacer, a property that may account for its ability to stimulate dnTA strongly despite an overall lower affinity for Cdc13.

### Limitations to the predictive capacity of the CATHI algorithm

The well-characterized and functional SiRTA 2R780(-) has a CATHI score of only 13, despite stimulating dnTA with an efficiency of 31.11% (Hoerr et al. 2023). By fluorescence polarization, this sequence competes for Cdc13 binding more effectively than many sequences with higher CATHI scores, suggesting that the failure of the algorithm to predict SiRTA function (at least in this case) is primarily a failure to predict Cdc13 binding. One possible explanation is that the CATHI algorithm does not prioritize matches to the GxGT motif identified as particularly impactful for Cdc13 binding (Anderson et al. 2002). Indeed, we find that mutation of even one GxGT motif in SiRTA 2R780(-) strongly reduces Cdc13 binding and nearly eliminates SiRTA function (Figure 7c) (Hoerr et al. 2023). The 226 bp minimal sequence of SiRTA 2R780(-) contains seven GxGT sequences that may account for the ability of this sequence to stimulate dnTA despite lacking sufficiently long/abundant telomere-like tracts to be identified by the algorithm.

Although the presence of GxGT motifs is an attractive explanation for the activity of SiRTA 2R780(-), attempts to incorporate the motif into the algorithm did not improve the accuracy with which SiRTA function could be predicted and instead increased the number of false positive results. For example, the false negative SiRTA 14R306(+) (CATHI score = 15.5) contains five GxGT motifs (two of which overlap), but the false positive SiRTA 12-330(+) (CATHI score = 22) also contains five distinct GxGT motifs. There are at least three (non-exclusive) explanations for remaining discrepancies between the predictive algorithm and measured rates of dnTA. First, it remains unclear how Cdc13 affinity is affected by deviation from the consensus telomere binding site. Although extensive mutagenesis has been conducted *in vitro*, these studies either altered single nucleotide sites (showing that positions 2 and 5-11 are tolerant of single changes) or simultaneously mutated the seven 3’-most nucleotides (showing that the GxGT motif is insufficient; Lewis et al. 2014; Glustrom et al. 2018). Neither approach fully recapitulates the sequences that Cdc13 will encounter at internal sites exposed by resection. Second, as described above, the distance between Cdc13 binding sites likely contributes strongly to SiRTA function. We have attempted to account for this property by using a window size of 75. However, some effects on SiRTA efficiency likely arise from varied distributions of Cdc13 binding sites that we cannot fully capture with the algorithm. Third, we suspect that dnTA can be stimulated either by a small number of high-affinity Cdc13 binding sites or by a larger number of low-affinity sites. SiRTAs at either extreme of this continuum may be difficult to identify using the current strategy.

### SiRTAs do not colocalize strongly with binding sites for other telomere/telomerase-associated proteins

Our co-localization analysis failed to identify additional proteins that strongly impact SiRTA function. Overall, we consider it unlikely that the observed enrichment represents a functional relationship. For example, Rap1 binding sites are overrepresented among SiRTAs, but this result is not surprising given that the consensus binding site for Rap1 is also TG-rich. In previous work, we showed that Rap1 association *per se* is not required for SiRTA activity (Obodo et al. 2016). Our results suggest that Est2 binding in undamaged conditions is not required for a sequence to function as a SiRTA. Only 8.1% of predicted SiRTAs overlap with experimentally determined sites of Est2 enrichment and only one of the fourteen active SiRTAs is also an internal Est2 binding site.

Consistent with our observation that SiRTAs stimulate dnTA even when located several kilobases from an induced DSB, we observe only a modest correlation between sequences that function as SiRTAs and sites enriched for phosphorylated H2A (γH2AX), a marker of DNA damage. Since enrichment was determined in undamaged cells, these sites represent regions of the genome that are prone to spontaneous damage, likely due to difficulties encountered during DNA replication. It will be interesting to determine whether SiRTAs that overlap with fragile sites are more likely to stimulate the spontaneous formation of gross chromosomal rearrangements. For example, we have proposed that the generation of acentric fragments through dnTA on chromosome II is facilitated by an unusually high density of inverted repeats in this region combined with errors in the resolution of stalled replication forks (Hoerr et al. 2023). In this light, it is interesting to note that sequences required to stimulate dnTA on chromosome II [SiRTA 2R780(-), coordinates 779784 to 780009] overlap with a region of enhanced γH2AX association (779987-780040; Capra et al. 2010).

Interestingly, we find a statistically significant tendency for predicted SiRTAs, if found within an open reading frame, to be oriented with the TG-rich sequence on the template strand. We propose that the presence of the TG-rich sequence within the exposed strand of the transcription bubble may be deleterious. Interestingly, this bias is opposite to that observed for G-quadruplex forming sequences in mammalian cells, which are more likely to be found on the coding strand (Agarwal et al. 2014; Rhodes and Lipps 2015; Kim 2019). In yeast, Replication protein A (RPA)-bound single-stranded DNA at stalled transcription complexes has been implicated as a major signal of DNA damage (Wang and Haber 2004; Tapias et al. 2004; Fanning 2006). Conceivably, competition for binding to single-stranded DNA by Cdc13 could interfere with this process, leading to selection against Cdc13 target sequences on the exposed coding strand. Despite the frequent presence of SiRTAs within genes, transcription does not appear to be required for SiRTA function since the test site that we developed on chromosome VII is contained within an intragenic region, more than 1.5 kilobases from the 3’ end of the *ADH4* gene.

### Implications of SiRTA distribution in the yeast genome

The compact and well-annotated yeast genome presents an opportunity to assess evidence of selective pressures that might act upon sequences with a propensity to stimulate dnTA. Based on the hypothesis that dnTA within the nonessential terminal region of a chromosome arm might provide a selective advantage by allowing a cell to survive a persistent DSB, we examined the distribution of predicted SiRTAs in essential and nonessential chromosome regions. As predicted, we found a significant enrichment of putative SiRTAs in nonessential terminal regions. Furthermore, as expected for a role in chromosome stabilization, only SiRTAs in the TG-orientation are overrepresented. However, both effects disappear when the subtelomeric X and Y’ elements are removed from the analysis (Figure 4a and c).

Nine of the TG-oriented SiRTAs in the subtelomeric regions correspond to stretches of TG_1-3_ (perfectly “telomeric”) sequence that are located predominantly between tandem Y’ elements. However, the vast majority, while TG-rich, deviate substantially from the TG_1-3_ pattern. Whether these sequences are vestiges of ancient telomeric repeats is unclear. Because the Y’ and X elements are similar between chromosome ends, many of the subtelomeric SiRTAs have similar or identical CATHI scores and represent a small number of unique sequences. Given the near ubiquity of TG-oriented SiRTAs within subtelomeric regions (identified at 31 of 32 chromosome ends), we speculate that these sequences are conserved, at least in part, due to an ability to stimulate dnTA in the event of catastrophic telomere loss, most likely due to replication errors within the telomeric repeats. The subtelomeric region on the right arm of chromosome VI contains a truncated X element followed immediately by TG_1-3_ telomeric repeats and therefore lacks sequences predicted to function as a SiRTA (Figure 3c). This subtelomeric structure may have resulted from a prior dnTA event within the X element. In the future, it would be interesting to determine whether the right arm of chromosome VI is more sensitive to catastrophic loss of telomeric repeats than other chromosome termini that contain intact X element repeats.

While the spatial distribution and orientation of putative SiRTAs outside the subtelomeres are not strongly skewed, the number of sequences with potential to act as SiRTAs is significantly higher than predicted by chance (Figure 5). This excess is observed at both low and high scores, but is increasingly pronounced at higher CATHI scores. Scores of 30 or greater are approximately 20-fold overrepresented in the yeast genome compared to the random expectation (Figure 5b). Our data do not provide evidence that SiRTA function *per se* is driving this excess, particularly because we also observe an excess of scores below 20, representing sequences that are not likely to stimulate dnTA at unusually high levels (Figure 5c). Since many SiRTAs are located within coding regions, we considered the possibility that codon bias might explain this pattern. However, codons consisting of only G and T (or only C and A; TTT and AAA excluded) collectively are not overrepresented among all codons (Supplementary File 10; Nakamura et al. 2000). It is possible that particular amino acid repeats could result in this effect. For example, poly-proline tracts consisting of CCA and CCC codons can generate a SiRTA signature. However, only a small fraction of poly-proline tracts are also identified as potential sites of dnTA.

An intriguing possibility is that association of Cdc13 with single-stranded DNA revealed by resection may be important to stimulate fill-in synthesis of the resected strand. At telomeres, Cdc13, in association with its binding partners Stn1 and Ten1, recruits polymerase α-primase to facilitate resynthesis of the 5’ recessed strand (Grandin 2001; Rice and Skordalakes 2016; Ge et al. 2020). In mammalian cells, the complex of Ctc1, Stn1, and Ten1 fulfills the same role at both telomeres and double-strand breaks (Chastain et al. 2016; Giraud-Panis et al. 2010; Wang et al. 2007). In this model, TG-rich sequences may be retained in the genome at a higher-than-expected frequency to facilitate DNA repair, with elevated rates of dnTA resulting as a rare byproduct.

Persistence of sequences such as the long TG-dinucleotide repeat on chromosome VI that support extremely high levels of dnTA is surprising since it seems likely that such sequences would interfere with normal repair. Future work will address whether dnTA is inhibited at SiRTAs in some contexts (for example, the SiRTA on chromosome VI may be less capable of stimulating dnTA in its endogenous location than when that same sequence is integrated at the test site on chromosome VII). Alternatively, the deleterious consequences incurred by dnTA at such a sequence may be insufficient to result in purifying selection or the TG-dinucleotide repeat may contribute to cell fitness through some other mechanism.

This work provides, for the first time, a genome-wide map of sites predicted to stimulate dnTA. With the exception of sites clustered in subtelomeric regions, the largely random distribution and orientation of SiRTAs throughout the genome stands in interesting contrast to the observation that sequences with this capability are found much more often than expected by chance. The tools presented here will facilitate studies to address this apparent contradiction and to determine the impact of these sequences on genome stability and evolution.

## Supporting information

Supplementary figures Ngo et al.

## Data availability

Strains and plasmids are available upon request. Code to run the CATHI algorithm can be found at https://github.com/bentonml/cathi. Code to model random distribution can be found at https://github.com/geofreyfriedman/sirta. Data summarized in Figures 1 and 2 are found in Supplementary File 3. Data summarized in Figure 3 are found in Supplementary Files 2 and 4. Data summarized in Figure 4 are found in Supplementary Files 2 and 5. Data summarized in Figure 5 are found in Supplementary File 6. Data summarized in Figure 6 are found in Supplementary Files 3 and 5. Data summarized in Figure 7 are found in Supplementary File 7. Locations of data presented in supplementary figures are referenced within the corresponding figure legends. Sequencing data are available from the NCBI Sequence Read Archive (SRA) under BioProject ID <u>PRJNA939836</u>.

## Acknowledgements

We thank James Haber and Deborah Wuttke for providing us with strains and plasmids. We also thank Katherine Paulin for her guidance regarding the protein binding experiments. We are grateful to the VANTAGE sequencing core for performing the Illumina sequencing in this work.

## Funding

This work was supported by National Institutes of Health award R01 GM123292 and a Vanderbilt-Ingram Cancer Center (VICC) Shared Resource Scholarship to K.L.F. E.A.H. is supported by the National Institutes of Health National Eye Institute (F31 EY033235). Research in A.R.’s lab is supported by grants from the National Science Foundation (DEB-2110404), the National Institutes of Health National Institute of Allergy and Infectious Diseases (R01 AI153356), and the Burroughs Wellcome Fund. Research in B.F.E’s lab is supported by NIH grant R35 GM136401. R.E.H. and D.I.G. are supported by the National Institutes of Health award T32 GM137793. MARC Scholars T.G. and S.P. are supported by a grant from the National Institute of General Medical Sciences of the National Institutes of Health: T34 GM136451. The content of this paper is solely the responsibility of the authors and does not necessarily represent the official views of the National Institutes of Health.

## Conflicts of Interest

A.R. is a scientific consultant for LifeMine Therapeutics, Inc.

## Notes

### Summary of Updates

Supplemental files added.

