## Supplementary figures Ngo et al. for "A comprehensive map of hotspots of de novo telomere addition in *Saccharomyces cerevisiae*"

Supplementary Figure 1

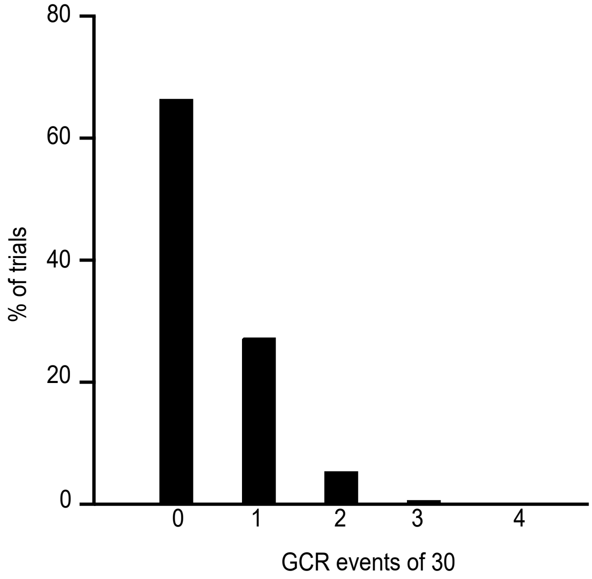


Supplementary Figure 1. Modeling the random distribution of GCR events within a 300 bp sequence inserted on chromosome VII. Given a 21,922 bp region between the HO cleavage site and the first essential gene on VII-L (*BRR6*), the number of randomly distributed events (of 30) expected to occur in the 300 bp sequence was determined over 10,000 trials. See text for explanation.

Supplementary Figure 2

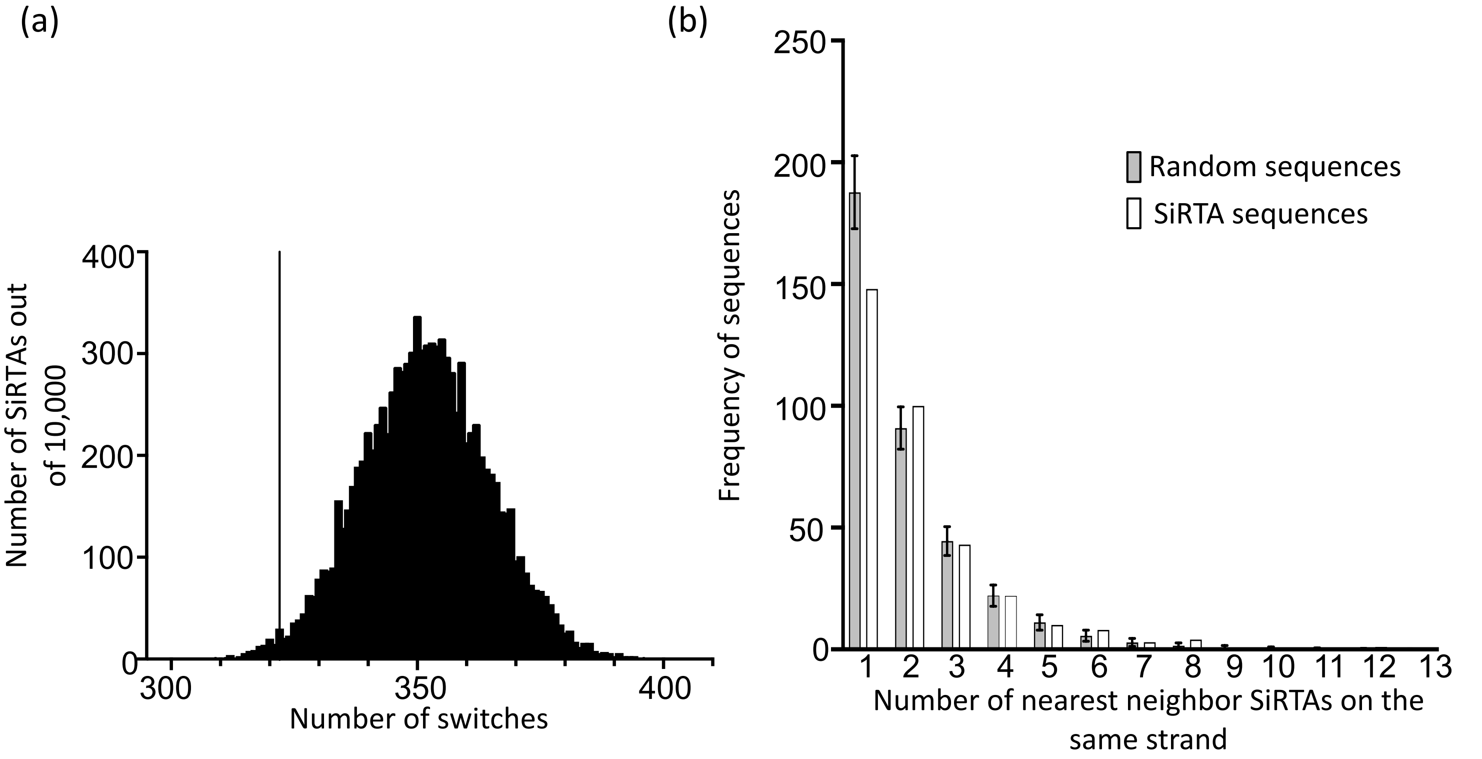


Supplementary Figure 2. Modeling the random strand distribution of SiRTAs. Using the known number of predicted SiRTAs on the top and bottom strand of each chromosome, as found in Supplementary File 4, 10,000 iterations were generated in which those SiRTAs were randomly distributed between strands. Distributions were analyzed in two ways. In (a), the number of times that neighboring SiRTAs were found on different strands (number of “strand switches”) was calculated. The observed value of 322 is shown (line), corresponding to a probability of 0.013. In (b), the number of adjacent SiRTAs on the same strand (the “run length”) was tabulated and summed across all chromosomes for each of 10,000 iterations. Average and standard deviation is shown for the randomized trials (grey bars). The actual values are shown with the white bars. Only cases in which both flanking SiRTAs are on the opposite strand (runs of one) are statistically underrepresented relative to expectation (p = 0.0043).

Supplementary Figure 3

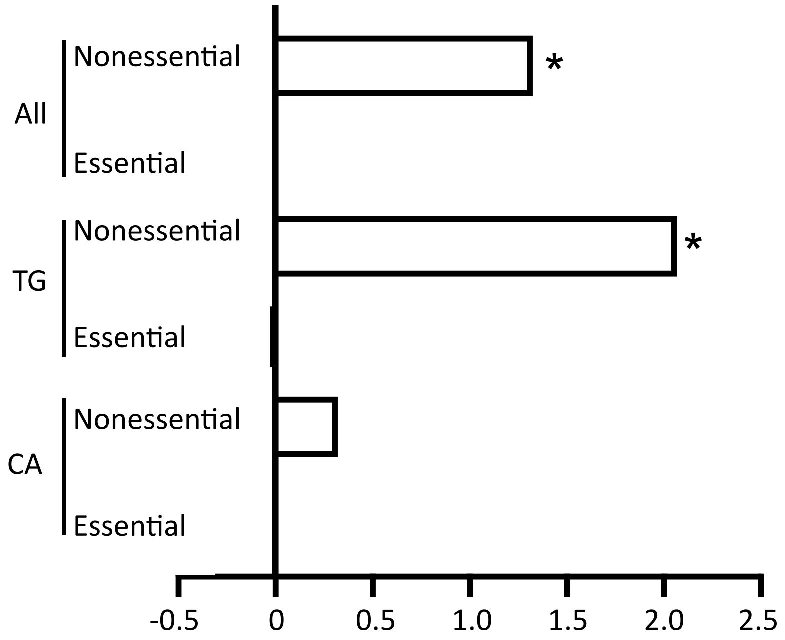


Supplementary Figure 3. SiRTAs are enriched in subtelomeric regions when excluding perfect telomeric repeats. Using a permutation strategy as described in Materials and Methods, the enrichment of SiRTAs (Log2 fold change) was determined for the nonessential and essential chromosome regions. This analysis is identical to that shown in Figures 4a and 4c except that nine subtelomeric SiRTAs containing perfect telomeric repeats (Supplementary File 5) were excluded. Analysis utilized all genomic sequences (except the terminal telomeric repeats). Essential and nonessential regions are defined in Supplementary File 2. *p-value <0.01 by chi-squared test with Bonferroni's correction.

Supplementary Figure 4

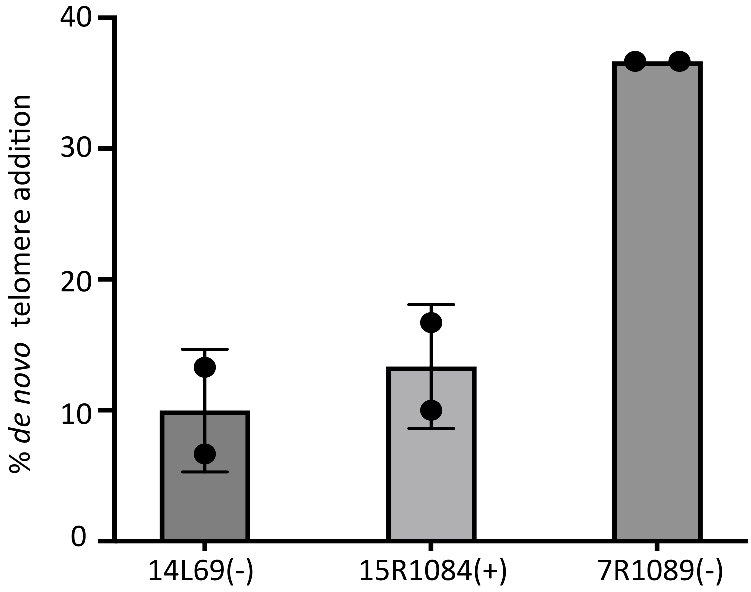


Supplementary Figure 4. Representative predicted SiRTAs in the X and Y' elements stimulate de novo telomere addition. The indicated SiRTAs (300 bp) were integrated at the test site on chromosome VII. Each data point represents one PCR experiment in which 30 GCR events were analyzed as described in Materials and Methods. Average and standard deviation are shown. SiRTAs 14L69(-) and 15R1084(+) are contained within X elements while 7R1089(-) is found within a Y' element. This data is summarized in Supplementary File 3.

Supplementary Figure 5

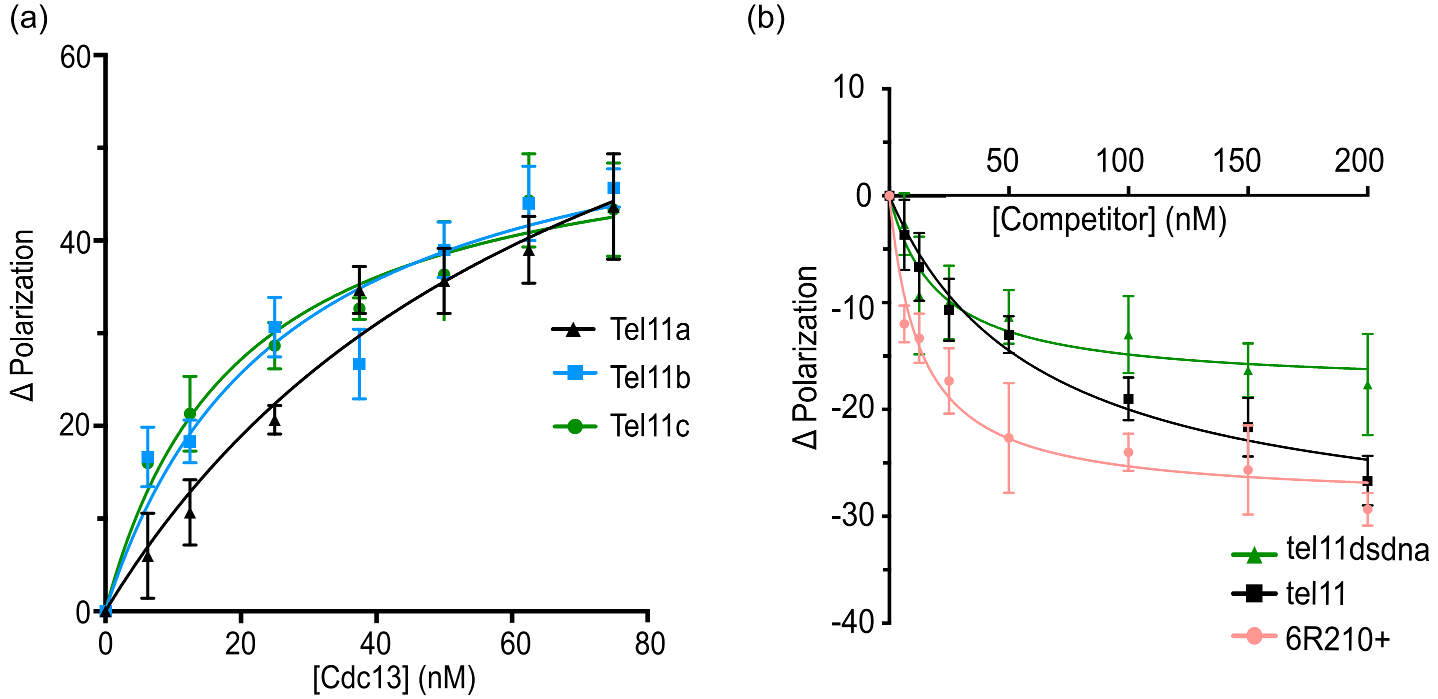


Supplementary Figure 5. Association of Cdc13 DNA binding domain (Cdc13-DBD) with DNA substrates. a) Each fitted curve represents a separate experiment to detect binding of Cdc13-DBD to the labeled Tel11 sequence. Individual data points are the average of three technical replicates; error bars are standard deviation. b) Representative competition experiment conducted as described in Materials and Methods. Tel11 is the labeled oligonucleotide. Results are shown for three different unlabeled competitor oligonucleotides (see Supplementary File 1 for sequences). Individual data points are the average of three technical replicates; error bars are standard deviation. Data summarized in Supplementary File 7.

Supplementary Figure 6

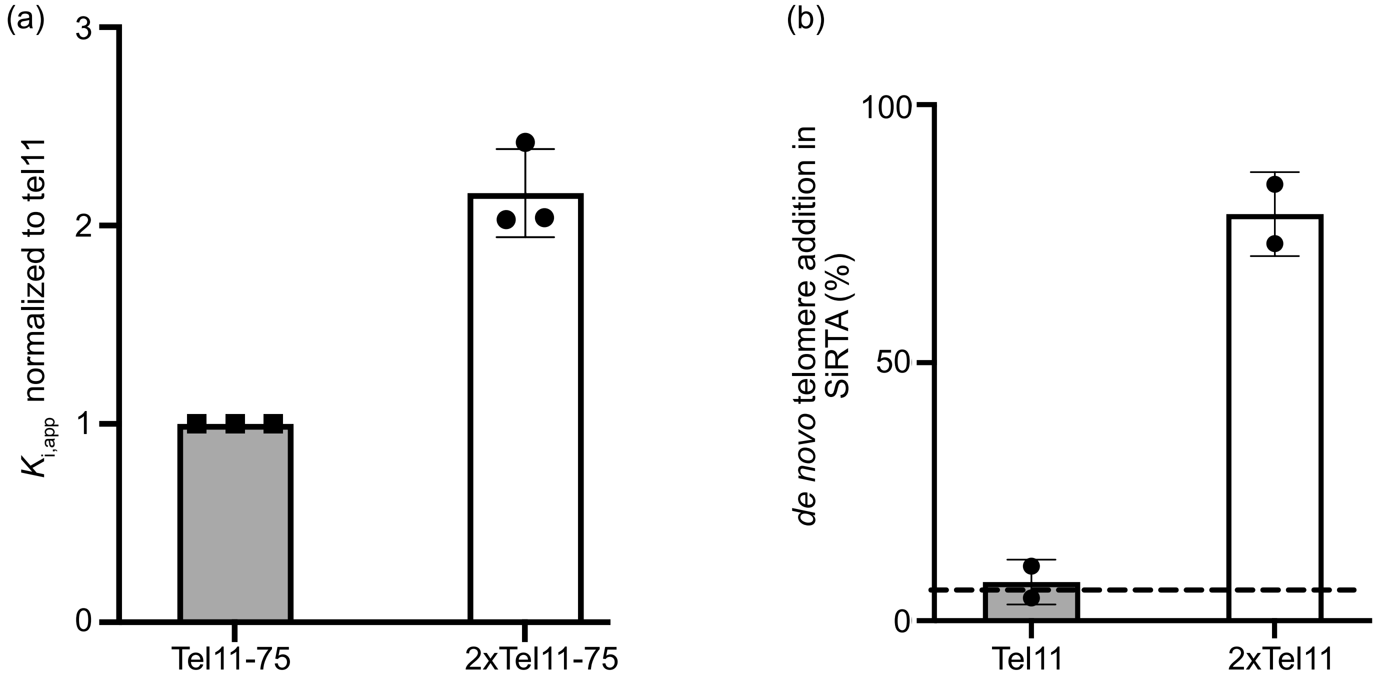


Supplementary Figure 6. Integration of two canonical Cdc13 binding sites is sufficient to stimulate high levels of dnTA. a) A competition fluorescence polarization assay was utilized to measure the relative association of the Cdc13 DNA binding domain with the indicated sequences (Supplementary File 1c). Relative $K_{i,app}$ was determined as described in Materials and Methods. Each point represents an independent measurement; error bars are standard deviation. Data for this figure is found in Supplementary File 7. b) The percent of GCR events involving dnTA within the indicated sequence was determined by PT-seq on chromosome VII. Tel11 contains a single canonical Cdc13 binding site; 2xTel11 contains two tandem sites. The dotted line represents the 6.6% threshold used to classify a sequence as a SiRTA. Data summarized in Supplementary File 3.

Supplementary Figure 7

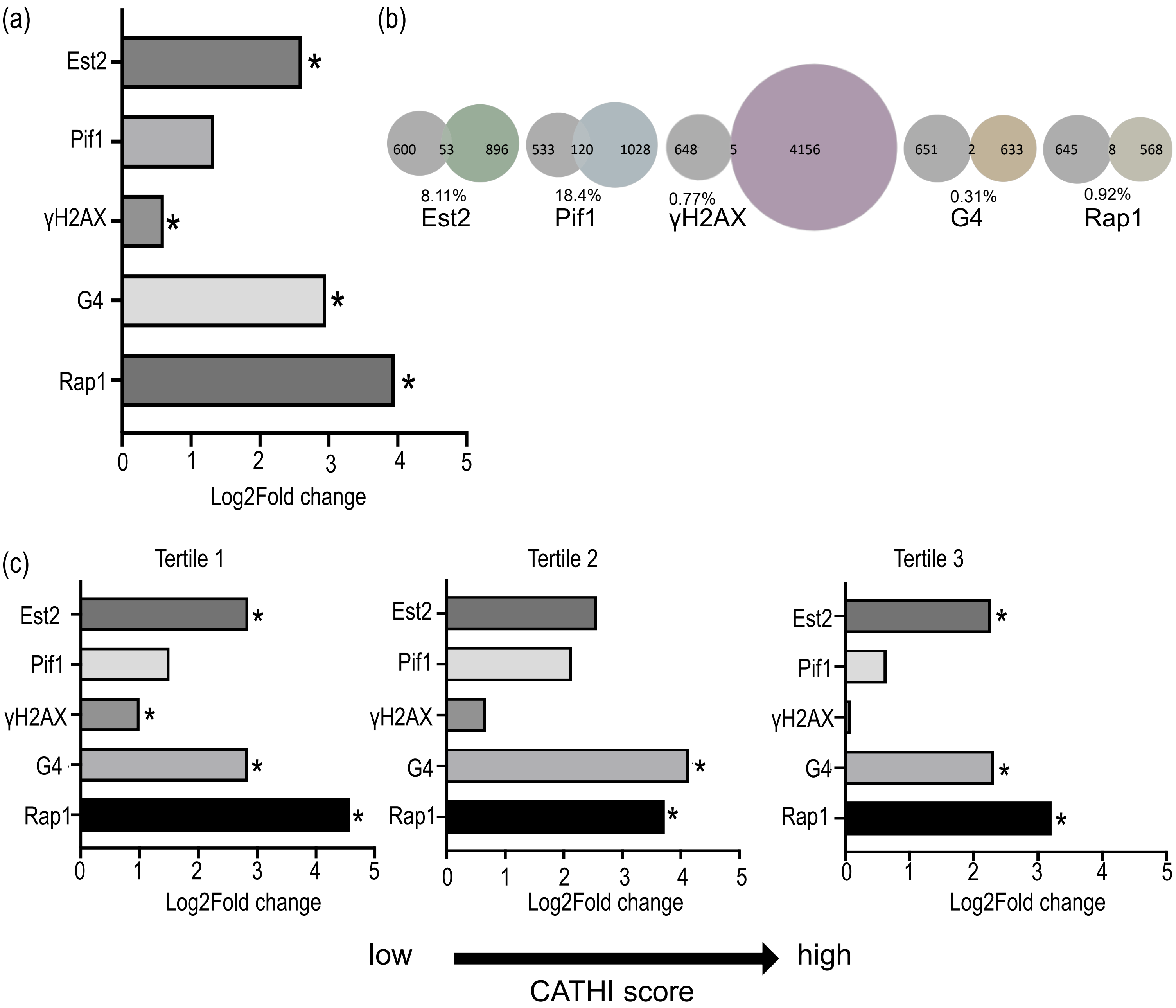


Supplementary Figure 7. Overlap of SiRTAs with protein binding sites and G-quadruplex forming sequences. a) Using a permutation strategy, the enrichment of SiRTAs (Log2 fold change) for overlap with the indicated protein binding sites and with G-quadruplex forming sequences was determined (see Materials and Methods). Analysis excluded sub-telomeric regions. *p-value<0.01 using Bonferroni's correction. b) Venn diagrams showing the number of SiRTAs that overlap with the indicated protein binding site or chromosome feature. Data used to make this figure is from Supplementary File 8. c) As in (a), except that SiRTAs were divided into tertiles based on CATHI score prior to permutation analysis. Tertile 1 represents the lowest scores. *p-value<0.01 using Bonferroni's correction.

Supplementary Figure 8

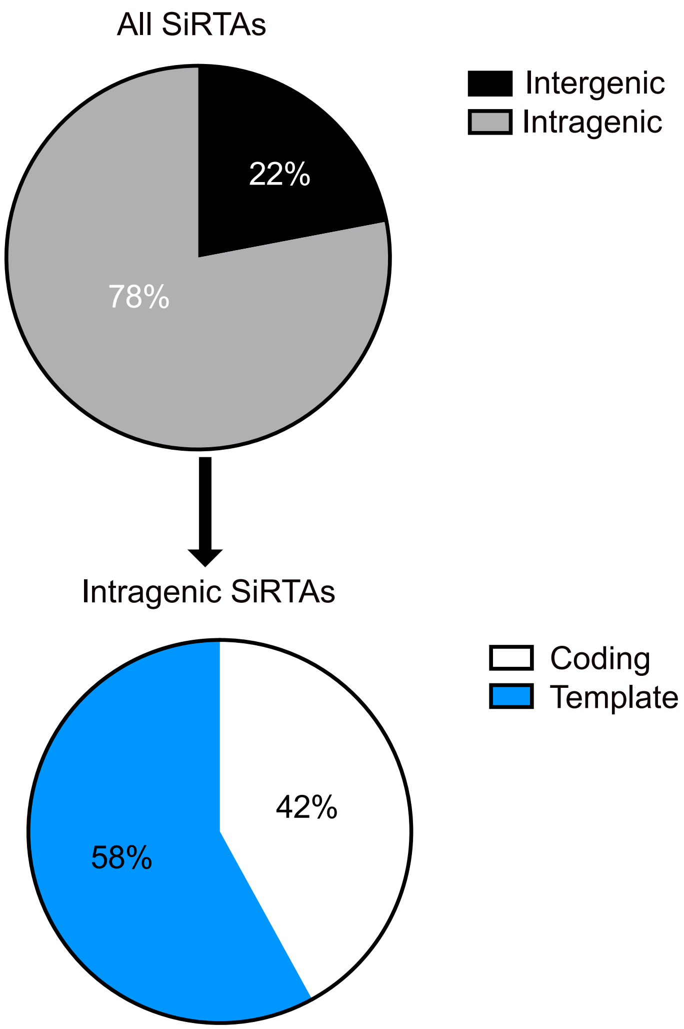


Supplementary Figure 8. Analysis of SiRTA overlap with protein coding regions. SiRTAs were classified as intragenic or intergenic (see Materials and Methods and Supplementary File 9). Intragenic SiRTAs were analyzed to determine if the TG-rich SiRTA sequence is located on the template or coding strand.
